# Multi-site enzymes as a mechanism for bistability in reaction networks

**DOI:** 10.1101/2021.05.06.442945

**Authors:** Clarmyra Hayes, Elisenda Feliu, Orkun S Soyer

**Author notes:** **Correspondings Authors**: Elisenda Feliu, Department of Mathematics, University of Copenhagen, DK-2100 Copenhagen, Denmark. Orkun S Soyer, University of Warwick, Coventry, CV4 7AL, UK. **Authors contributions**: CH and OSS have devised the study. CH, OSS, and EF performed analyses and simulations, interpreted the results, and wrote the manuscript.

## Abstract

Here, we focus on a common class of enzymes that have multiple substrate-binding sites (multi-site enzymes), and analyse their capacity to generate bistable dynamics in the reaction systems that they are embedded in. Using mathematical techniques, we show that the inherent binding and catalysis reactions arising from multiple substrate-enzyme complexes creates a potential for bistable dynamics in a reaction system. We construct a generic model of an enzyme with *n* substrate binding sites and derive an analytical solution for the steady state concentration of all enzyme-substrate complexes. By studying these expressions, we obtain a mechanistic understanding for bistability and derive parameter combinations that guarantee bistability and show how changing specific enzyme kinetic parameters and enzyme levels can lead to bistability in reaction systems involving mjulti-site enzymes. Thus, the presented findings provide a biochemical and mathematical basis for predicting and engineering bistability in multi-site enzymes.

## INTRODUCTION

Cellular reaction networks enable cells to remain out of thermodynamic equilibrium and to respond to external cues. The dynamics of these networks enable cellular homeostasis and decision making (1,2). Many decision-making processes involve so-called bistable dynamics, in which a system can attain two different steady states depending on initial conditions. Bistability is implicated in many cellular decision processes, including the cell cycle control (3), lysis-lysogeny decision (4), metabolic shifting (5-7), and persister formation (8).

Manifestation of bistability requires some mechanism of feedback (9, 10). In the case of enzymatic reaction systems, feedback dynamics can arise from transcriptional, or substrate- or product-based regulation, or via post-translational modification of enzymes. Several models implementing these types of enzyme regulation are shown to display bistability and are used to explain different cellular responses (2, 5, 6, 11-13). In the case of substrate- and product-based regulation of enzymes, a commonly used model considers an enzyme with two binding sites, where binding of substrate at one side leads to catalysis, while binding of the substrate or product on the other site alters catalytic rate. In such a two-site enzyme model, both bistability and oscillations are attainable depending on the specific binding mechanisms and the assumed functional forms of the rate equations (2, 11, 14-15). Despite this wide application of the two-site enzyme model, it is currently not clear how exactly a multi-site enzyme facilitates bistability and under which parameter regions and biochemical conditions it does so. This is a relevant question, considering that many enzymes found in central metabolism and signalling pathways are multimers comprising multiple substrate binding sites (16). Specific examples include dehydrogenases with key metabolic substrates (e.g. phosphoglycerate, malate and lactate) and commonly composed of dimers or tetramers with multiple binding sites (17), and kinases such as phosphofructokinase, which have multiple active binding sites (18). A better understanding of reaction dynamics of multi-site enzymes can allow us to predict which naturally existing enzymes might be implementing bistability for cellular decision making or might be suited for engineering of bistability through synthetic biology.

Here, we undertake an extensive theoretical study of a generalised model of an enzyme with *n* substrate binding sites, in order to derive both a biochemical intuition and a set of mathematical conditions on kinetic parameters for bistability. We use primarily analytical approaches to show that the multi-site nature of an enzyme inherently results in a potential for bistability. We then use this insight to derive conditions on the kinetic rate parameters of simple reaction networks with multi-site enzymes, that guarantee bistability for some concentration of substrate and enzyme. These findings allow us to predict and outline enzyme engineering strategies that can be employed to achieve bistability in simple reaction networks.

## RESULTS

To better understand how a multi-site enzyme can lead to bistability, we first create a generic model of substrate (*S*) to product (*P*) conversion mediated by an enzyme (*E*) that has *n*-substrate binding sites (Fig. 1A). In this initial model, we assume that the total concentration of substrate and product, and the total concentration of free and substrate-bound enzyme are conserved (see *Methods* and *Supplementary Information* (*SI*)). The former assumption is directly applicable when the substrate is a conserved moiety, such as enzyme co-factors or energy and reducing power equivalents (e.g. ATP-ADP and NADH-NAD^+^ pairs) (2, 19). This assumption is useful to illustrate our results, and relaxing it – as discussed below - show that our main conclusions remain intact for the cases where substrate concentration is freely changing (e.g. through fluxes by other reactions). The latter assumption of total enzyme concentration being conserved reflects the fact that the time scales of enzyme expression are in most cases slower compared to reaction dynamics.

**Figure 1.**
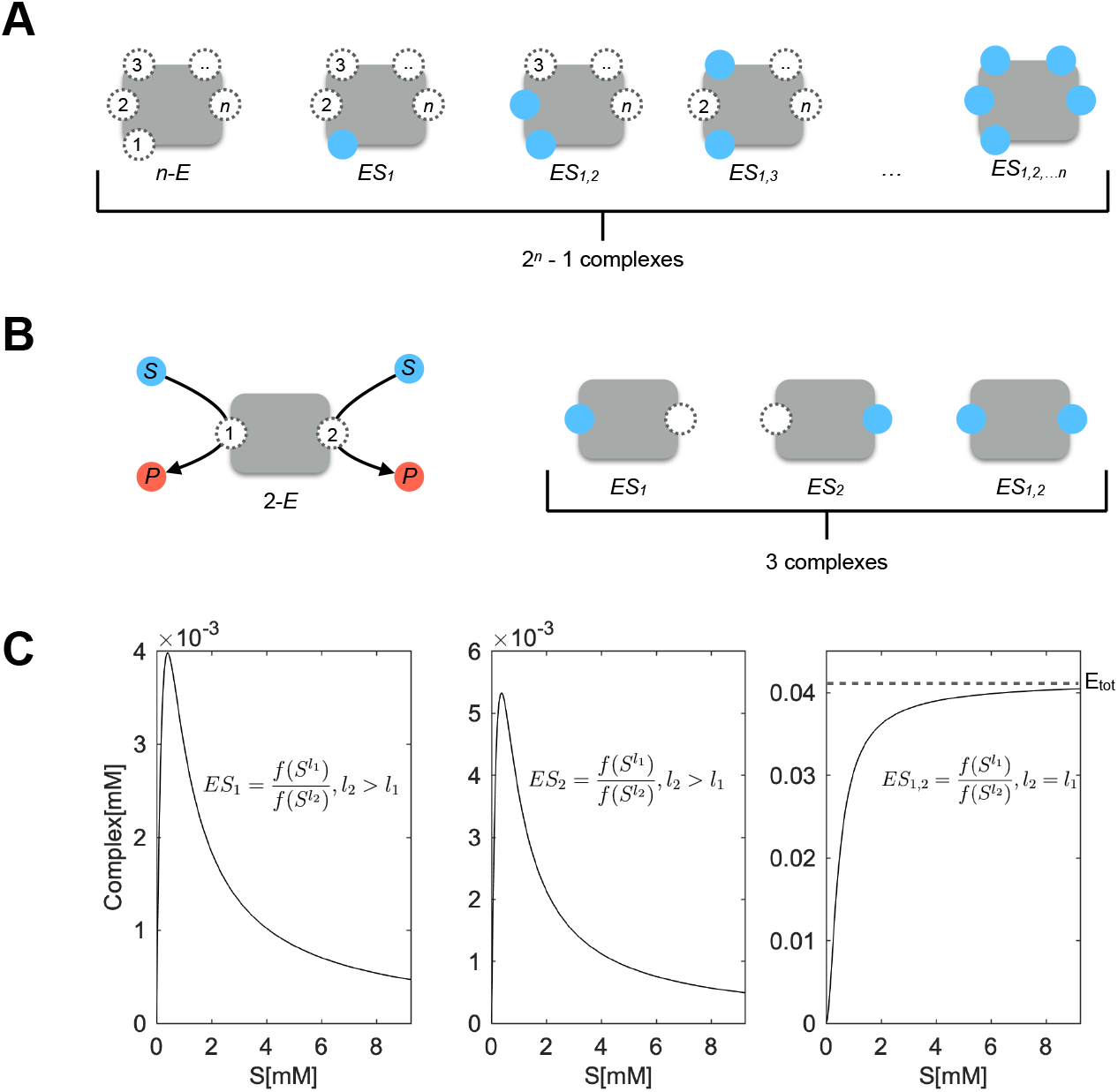
**A**. Cartoon representation of a generic *n*-site model, where *n*-E indicates an enzyme with *n* substrate binding sites. The substrate binding sites are numbered in a consecutive fashion and substrate-bound sites are shown in blue. Note that there are 2*n* – 1 possible substrate-enzyme complexes. **B**. Cartoon representation of a 2-site enzyme model. The substrate (*S*) and product (*P*) are shown in blue and red respectively. Substrate binding is allowed in any order on each site, and both sites are assumed to have catalytic activity. The 3 possible substrate-enzyme complexes are shown on the right. See *Methods* for reactions and differential equations for this 2-site enzyme model. **C**. The steady state concentration of each of the substrate-enzyme complexes with increasing concentration of substrate. The parameters, as listed in Eq. 4, are set to the following values for these simulations; *k*_*1*_ = *k*_*4*_ = *k*_*6*_ = *k*_*10*_ = 10^8^ M^-1^min^-1^, *k*_*2*_ = *k*_*5*_ = *k*_*7*_ = *k*_*11*_ = 10^4^ min^-1^, *k*_*3*_ = 10^5^, *k*_*12*_ = 1.5 10^5^ min^-1^, *k*_*8*_ = *k*_*13*_ = 10^3^ min^-1^, *S*_*tot*_= 2.31 10^−3^ M, *E*_*tot*_ = 4.15 10^−5^ M. Panels from left to right show the steady state concentrations of the two single-substrate complexes, and the fully-bound complex. A simplified version of Eq. 2, describing the steady state concentration of the complexes is shown on each panel, highlighting the degree of the polynomials. On the right-most panel, the dashed line indicates total enzyme concentration.

To make the model as generic as possible, we use mass-action kinetics with irreversible enzymatic catalysis, and consider substrate molecules binding to the enzyme in any order and also irrespective of how many substrates are already bound. As we show in the *SI*, more restricted assumptions about substrate binding order or affinity, do not alter our main conclusions. To exemplify our modelling approach, in the *Methods* section, we provide the set of reactions arising from the generic model for a 2-site enzyme, i.e. *n* = 2 (see also Fig. 1B). For our general *n*-site model, the full set of reactions can be formally written as:

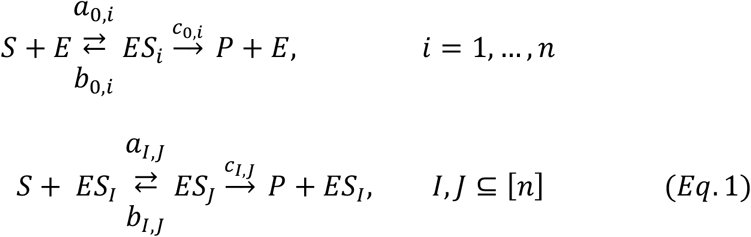

where *a, b* and *c* are kinetic rate constants associated with the individual substrate binding sites *i*, which are numbered 1 through *n*. The set [*n*] is the complete set of binding sites [*n*]={1, …, *n*}, and *ES*_*I*_ and *ES*_*J*_ are enzyme complexes in which a given number of substrate molecules are bound respectively to a set of sites *I* and *J*. In other words, *I* and *J* are sets with *any* number of elements from the list of sites 1 through to *n;* (*I, J* ⊆ [*n*]). For example, for *I* = {1,3,4}, *ES*_*I*_ is the enzyme complex where the sites numbered 1, 3, 4 are bound to substrate molecules (Fig. 1A). Additionally, *ES*_*J*_ is formed by the binding of a single, additional substrate molecule to *ES*_*I*_, meaning the difference between the sets of *I* and *J* in Eq. 1 is one element. Note also that the system defined by Eq. 1, results in 2^*n*^-1 enzyme complexes (Fig. 1A).

### Fully-bound and non-fully-bound enzyme complexes display distinct steady state dynamics with increasing substrate concentration

We analysed the above generic model using analytical methods to derive solutions for the steady state concentrations of all 2^*n*^ – 1 enzyme complexes, as functions of the steady state concentration of substrate ([*S*]) (see *SI* for details). We found that the steady state concentration of any complex (*ES*_*I*_) is given by:

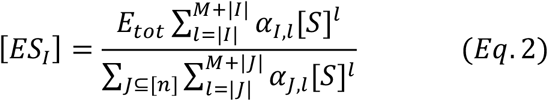

Here, *E*_*tot*_ is the total enzyme concentration including both free and bound forms of the enzyme, and *M* = 2^*n*^ – 1 – *n*. The terms |*I*| and |*J*| are the number of elements (i.e bound sites) in a given complex, and thus, the index *l*, which also appears as an exponent to [*S*], is over the number of bound substrates. The terms *α*_*I,l*_ and *α*_*J,l*_ indicate a positive function of the kinetic reaction constants associated with each of the enzyme complexes (see *SI* for details and *Methods* for an example with *n* = 2). We note that Eq. 2 is derived under the most generic case of substrates binding to different enzyme sites in any order, however, we show that Eq. 2 remains true if we assume more specific binding processes, e.g. binding at a specific enzyme site requiring other sites to be bound with substrate (see *SI*, Section 1.1 for details). In such cases, some of *α*_*I,l*_ and*α*_*J,l*_ might be zero.

A close inspection of Eq. 2 shows that [*ES*_*I*_] will always be given by a fraction of two polynomials in [*S*]. These polynomials will differ in their degree in [*S*] unless *I* is equal to the full set (i.e. *I*=[n]). This is because, when *I≠*[n], the summations in the denominator and the numerator in Eq. 2 are over different numbers of bound substrates. Specifically, the summation in the denominator is over all enzyme complexes and the largest degree of this polynomial will be equal to *M* + *n =* 2^*n*^ – 1, the total number of enzyme complexes. In contrast, the summation in the numerator is over the enzyme complexes that can be generated from the enzyme complex *ES*_*I*_. If *ES*_*I*_ is the fully-bound enzyme complex, then the degree of the numerator will be equal to that of the denominator, as the largest possible value of the index *l* would be *M* + | *I* | *=* 2^*n*^ – 1. If *ES*_*I*_ is not the fully-bound complex, then the degree of the numerator will be equal to that of the denominator minus the number of empty binding sites in *ES*_*I*_. For instance, if the enzyme has two substrate binding sites, leading to three potential different enzyme complexes, the degree of the polynomial in the denominator will be three (Fig. 1B and C). The polynomial in the numerator would have a degree of three for the fully bound complex, while for the two complexes, consisting of one filled and one empty binding site, it would have a degree of two (Fig. 1B and C).

The specific structure of Eq. 2 provides an insight to the behaviour of steady state concentrations of the different enzyme complexes with increasing [*S*] (Fig. 1C). Considering the fact that the polynomials comprising Eq. 2 have positive coefficients (given by *α*_*I,l*_ and *α*_*J,l*_, which are functions of kinetic rates), the steady state concentration of all enzyme complexes will initially increase from zero with increasing [*S*]. Since Eq. 2 for the fully-bound complex has polynomials of the same degree in the numerator and denominator, the limit value of Eq. 2 at very high [*S*] for this complex will be the ratio of the coefficients of the highest degree terms of the numerator and denominator. We show that this ratio is equal to *E*_*tot*_, the total enzyme concentration in the network (see *SI*, Theorem 1). Thus, for the fully-bound enzyme complex the steady state concentration will initially increase with increasing [*S*] and approach finally a positive value given by *E*_*tot*_ (Fig. 1C, last panel). In the case of the non-fully-bound enzyme complexes, Eq. 2 will have a lower degree polynomial in the numerator than the denominator, and therefore, its limit value at very high [*S*] will approach zero. Thus, for the non-fully-bound enzyme complexes their steady state concentration will initially increase with increasing [*S*], show at least one peak, and then approach towards zero from above (Fig. 1C, first two panels).

Note that, while Fig. 1C shows the behaviour of Eq. 2 for an enzyme with *n* = 2, the analytical summary presented here is independent of *n*. It shows that, for a multi-site enzyme, we will always have two distinct, and qualitatively different curves describing the different enzyme complexes’ steady state concentrations (as exemplified in Fig. 1C). From here on, we refer to these two qualitatively distinct types of curves as ‘positive’ and ‘negative’ type, respectively. Both positive and negative type curves will increase when [*S*] is small and increasing. At large values of [*S*], both curves will approach a limit value, with positive type curve approaching its limit from below and a negative type curve approaching its limit from above (Fig. 1C). These overall conclusions for curve shapes against small and large values of [*S*] are independent of the specific values of the kinetic rate parameters. They arise solely because of the polynomial degree structure of Eq. 2, in other words, from the multi-site structure of the enzyme.

The exact shape of the curves for intermediate, increasing values of [*S*], however, and in particular the number of peaks they will display before approaching the limit value, will depend on the catalytic and Michaelis-Menten (*K*_*m*_’s) rate constants of the individual enzyme-substrate complexes (i.e the functions *α*_*I,l*_ and *α*_*J,l*_ in Eq. 2). For an enzyme with *n* = 2, the negative type curves (of the single substrate complexes) will always show a single peak and have one inflection point (see *SI*, Section 1.1). The positive type curve (of the fully-bound, two substrate complex) mostly shows no peaks and is a steady increasing function of [*S*], but there are kinetic parameters for which it would display peaks, as we discuss below (see *SI*, Section 2.4). With higher *n*, both the negative and positive type curves can readily display multiple peaks. Intuitively, and from a biochemical perspective, the positive type curve can be thought of as a saturation process, in which increasing [*S*] pushes more enzyme binding sites to be filled, ultimately leading to an increase of the steady state concentration of the fully-bound enzyme complex. Correspondingly, the steady state concentrations of the non-fully-bound enzyme complexes decrease with increasing [*S*], giving rise to the negative type curve.

### The negative type curves of non-fully-bound enzyme complexes underpin the potential for bistability

We now consider the catalytic flux through each enzyme complex. We refer to the catalytic flux through complex *ES*_*I*_, as, 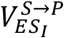 and note that its steady state value will be a function of the steady state complex concentration, [*ES*_*I*_]. Furthermore, the total catalytic flux through the enzyme, *V*_*S→P*_, will be given by the sum of the individual fluxes through each of its complexes. By the analysis above, *V*_*S→P*_ tends to *E*_*tot*_, times the sum of the catalytic rate constants of the fully-bound complex. To illustrate the ideas for general *n*, we consider first the example case for an enzyme with *n* = 2 (shown in Fig. 1B and 1C). It is easier to graphically understand how bistability arises in this system if we analyse the behaviour of the catalytic fluxes against [*S*_*sum*_] = *S*_*tot*_ - [*P*], where *S*_*tot*_ is a constant describing the combined amount of product and free and bound substrate (see *SI*, Section 1.1). As we show in the *SI*, for *n* = 2, [*S*_*sum*_] is an increasing function of [*S*] and hence, the qualitative behaviour of *V*_*S→P*_ against increasing [*S*] or [*S*_*sum*_] is the same.

In Fig. 2, we show 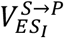 against [*S*_*sum*_] for two different parameter sets, and as expected, we see that the behaviour of 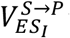 against [*S*_*sum*_] qualitatively follows that of [*ES*_*I*_] against [*S*] as given by Eq. 2 and shown in Fig. 1. In the example shown in Fig. 2A, where we have the same parameters as in Fig. 1C, the total catalytic flux *V*_*S→P*_ is dominated by the fluxes through the non-fully-bound complexes, and as such, *V*_*S→P*_ displays a negative type behaviour in [*S*_*sum*_]. In Fig. 2B, we see the results for a second set of parameters, where *V*_*S→P*_ is dominated by the flux through the fully-bound complex, and as a result, it displays a positive type curve in [*S*_*sum*_]. As illustrated by these examples, which type of behaviour *V*_*S→P*_ displays will depend on the specific values of the catalytic and Michaelis-Menten (*K*_*m*_’s) rate constants of the individual enzyme complexes.

**Figure 2.**
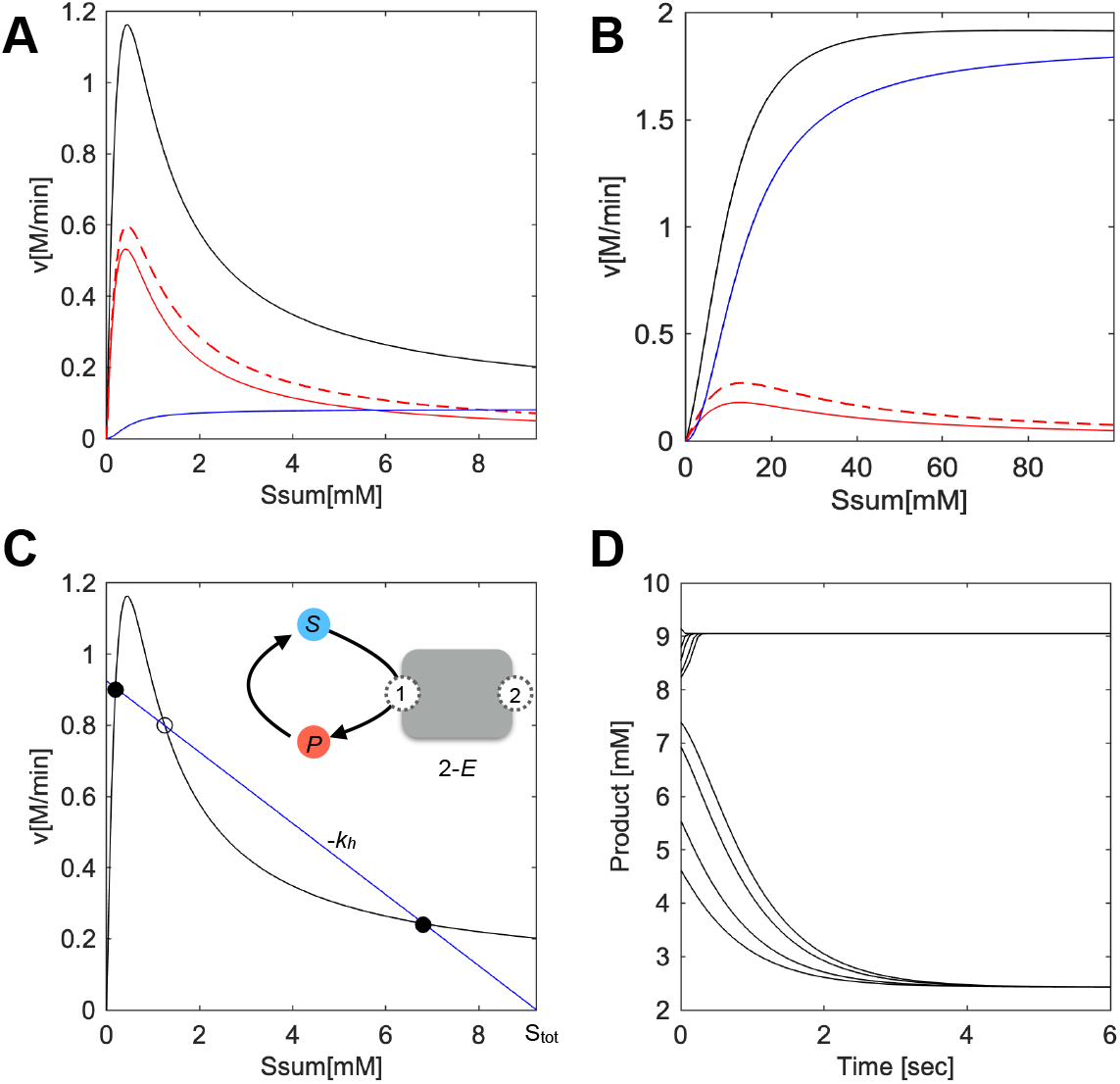
**A**. Steady state reaction flux through different substrate-enzyme complexes in the 2-site model against total substrate concentration. The straight and dashed red curves are for the reaction flux through the single-substrate bound complexes, while the blue curve is for the reaction flux through the fully-bound enzyme complex. The black curve shows the total reaction flux, i.e. *V*_*S→P*_. The parameters used are as in Fig. 1C. **B**. Steady state reaction flux through different substrate-enzyme complexes in the 2-site model against total substrate concentration. Curve shapes and colours have the same meaning as in part A. The parameters used are: *k*_*1*_ = *k*_*4*_ = *k*_*6*_ = *k*_*10*_ = 10^8^ M^-1^min^-1^, *k*_*2*_ = *k*_*5*_ = 10^4^ min^-1^, *k*_*3*_ = 10^5^, *k*_*7*_ = *k*_*11*_ = 10^5^ min^-1^, *k*_*8*_ = 2.5 10^4^ min^-1^, *k*_*12*_ = 1.5 10^5^ min^-1^, *k*_*13*_ = 2 10^4^ min^-1^, *S*_*tot*_ = 2.5 10^−2^ M, *E*_*tot*_ = 4.15 · 10^−5^ M. **C**. The 2-site enzyme embedded in a simple reaction system involving a back reaction from product to substrate, as shown on inset. The black curve shows the total reaction flux *V*_*S→P*_. The blue line shows the back reaction flux, i.e *V*_*P→S*_. Note that the intersection points of these two curves represent the steady state points in the system. These points are marked on the plot, with stable and unstable steady states represented with filled and open circles respectively. The parameters are the same as those used in Fig. 1C A, with *k*_*h*_ = 10^2^ min^-1^. **D**. Product concentration over time, resulting from a numerical simulation of the system shown in part C and using the same kinetic parameters as used there. Each curve shows the result of an individual numerical simulation, starting from a different initial condition.

We now consider the shape of the *V*_*S→P*_ curve in the context of a reaction system. To start with, we consider a simple scenario, involving a back reaction from product to substrate, creating a reaction cycle (see Fig. 2C). We initially assume that the product to substrate conversion is a non-enzymatic, hydrolysis type reaction, governed by a constant *k*_*h*_ (note that, below and in the *SI*, we relax this assumption without loss of the presented conclusions). The catalytic flux of this back reaction, *V*_*P→S*_, is given by *k*_*h*_ ·[*P*] and therefore, behaves linearly with increasing [*S*_*sum*_]. This linear relation has slope -*k*_*h*_ and intercept *S*_*tot*_ (Fig. 2C). When we plot *V*_*S→P*_ and *V*_*P→S*_ against [*S*_*sum*_] on the same plot, the intersection points represent the steady states of the reaction system, i.e. points where the product formation flux, *V*_*S→P*_, equals that of product loss, *V*_*P→S*_. Using the fact that *V*_*P→S*_ is a line with negative slope, we can see that a negative type *V*_*S→P*_ curve opens the possibility to have three intersections between *V*_*S→P*_ and *V*_*P→S*_, and therefore three steady states. Three steady states are the hallmark of bistability, and indeed, for this parameter set, our model displays bistability, where different starting conditions can lead to different steady state dynamics (Fig. 2D). Since adjusting the value of *S*_*tot*_ results in shifting the *V*_*P→S*_ line along the x-axis, we can graphically see that as long as *k*_*h*_ is below a certain threshold value, there will be some value of *S*_*tot*_ that ensures three intersections. In other words, tuning the *S*_*tot*_ value would allow shifting the *V*_*P→S*_ line across the x-axis on Fig. 2C, until three intersections with the *V*_*S→P*_ curve are obtained.

While we analyse a system with *n* = 2 and a sample parameter set in Fig. 2, we can use the above discussion to draw a general conclusion that will be true for any *n*. If *V*_*S→P*_ is of the negative type and its slope at the inflection point is smaller than the slope of *V*_*P→S*_ (that is -*k*_*h*_), then the curves will intersect three times if the line *V*_*P→S*_ passes through the inflection point. The slope of *V*_*S→P*_ at the inflection point depends on *E*_*tot*_ and the reaction rate constants, while the slope of *V*_*P→S*_ at the inflection point depends on *k*_*h*_. This graphical analysis, therefore, provides an intuition about why having a *V*_*S→P*_ of the negative type and with a slope at its inflection point smaller than -*k*_*h*_ provides a route to bistability in a system with a multi-site enzyme for some value of *S*_*tot*_. On the contrary, if *V*_*S→P*_ is of the positive type and does not display any peaks (as shown in Fig. 2B), bistability is precluded as *V*_*P→S*_ cannot intersect *V*_*S→P*_ in more than one point. When *V*_*S→P*_ is of the positive type and displaying a single or multiple peak, there is again the possibility for three intersection points and bistability (see *SI*, Section 2.4). In summary, this graphical discussion shows that a negative type curve for *V*_*S→P*_ guarantees three steady states after appropriately choosing the other relevant parameters (e.g. *k*_*h*_ and *S*_*tot*_).

### Kinetic rate parameter conditions that guarantee multiple steady states in a reaction system with a multi-site enzyme

In order to formalise and generalise the graphical considerations made above, we take a mathematical approach to determining conditions on kinetic parameters that result in multiple steady states. The idea is to identify the conditions when *V*_*S→P*_ is of the negative type, that is, when it converges to its limiting value from above, and use these conditions to guarantee that *V*_*P→S*_ and *V*_*S→P*_ will intersect at multiple points.

We find that *V*_*S→P*_ is of the negative type exactly when the following condition holds (*SI*, Section 1.2):

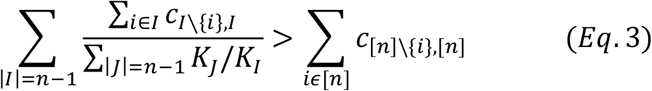

Here, *K*_*I*_ and *c*_*I*_\ {*i*},*I* represent the Michaelis Menten (*K*_*m*_) and catalytic rate constants as in Eq. 1, respectively, for the enzyme complexes with all binding sites bound but the *i*’th one (i.e. enzyme complexes with *n-1* sites bound). The term *c*_[*n*]_\ {*i*},[*n*] represents the catalytic rate constants of the fully-bound enzyme complex, where catalysis happens at the *i*-th binding site (see also Fig. 3).

**Figure 3.**
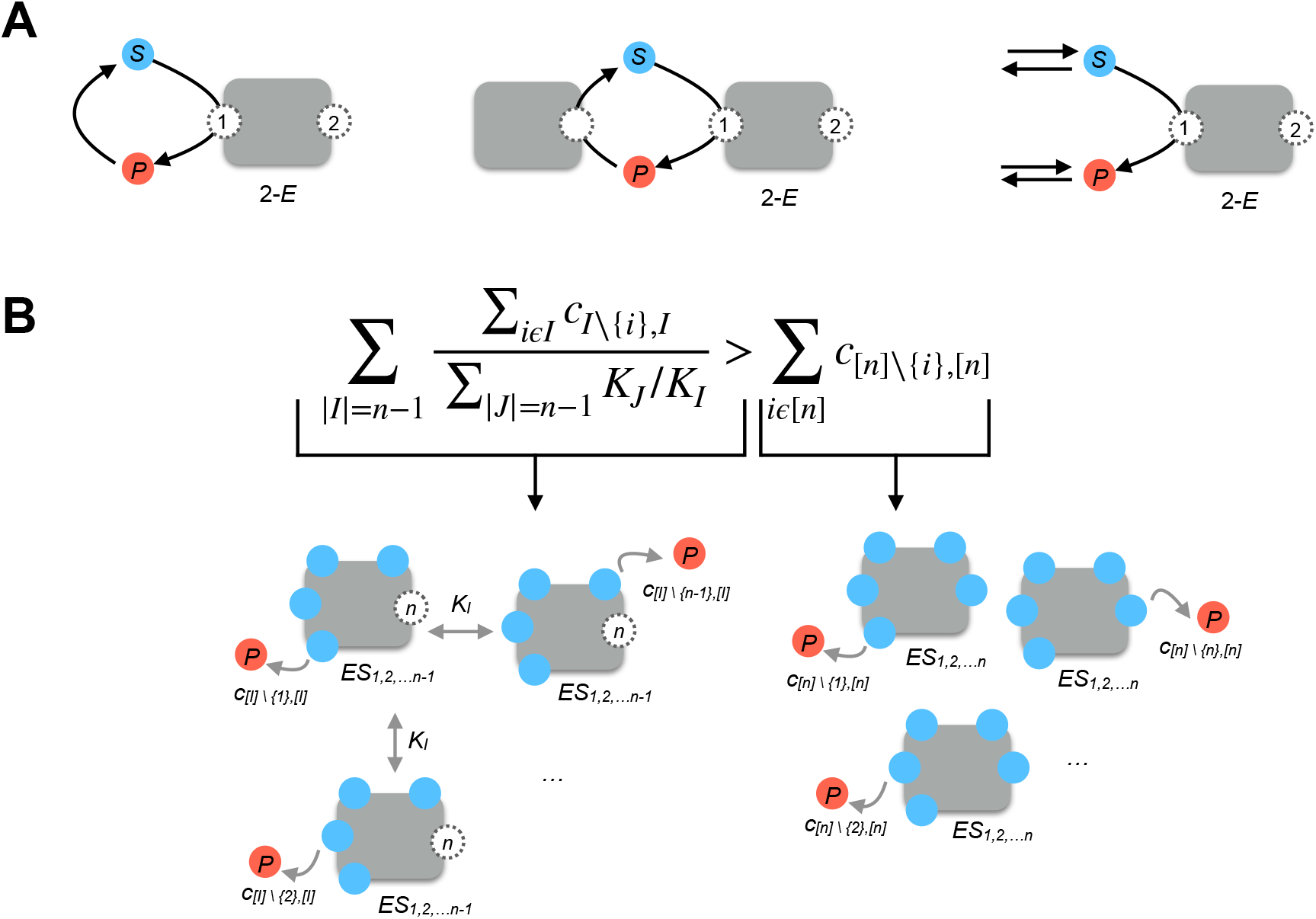
**A**. The three different reaction systems, embedding a multi-site enzyme, considered in this work. For simplicity, each system is shown with a 2-site enzyme model and with only a single reaction via one example binding site, while the mathematical analysis presented in the main text considers a *n*-site model with all possible binding and catalysis reactions. The resulting inequality for each 2-site system is provided under each cartoon, with the inequalities for the full model provided in the *SI*. **B**. The core inequality, as shown in Eq. 3 and common to all the cases considered, is written for the generic, *n*-site model. This inequality characterizes when *V*_*S→P*_ is of negative type. We note that the right side of this equation correspond to only the sum of catalytic rates from the fully bound enzyme complex, as depicted in the cartoon below. The right side of the inequality involves both catalytic rates and equilibrium constants of those enzyme complexes that are unbound only on one site.

We note that the condition defined by Eq. 3 is aligned with the graphical analyses we discussed in the previous sections (Fig. 1 and 2). There, we have shown that the curve type of *V*_*S→P*_ is determined by whether the fully-bound or non-fully-bound enzyme complexes are dominating the dynamics of catalysis. In line with these arguments, for Eq. 3 to hold and hence for *V*_*S→P*_ to be of the negative type, the sum of the catalytic rate constants for the *n-1* non-fully-bound complexes, each adjusted by the contribution of that complex in the system dynamics (represented by their *K*_*m*_’s), have to be greater than the sum of the catalytic rate constants of the fully-bound complex.

Eq. 3 determines the condition for *V*_*S→P*_ to be of the negative type. How this leads to multistability relates to the system, in which the multi-site enzyme is embedded in. We first study the simple case of a non-enzymatic back reaction from product to substrate (see next section for results of alternative reaction systems). We find that we are guaranteed to have three positive steady states in such a cyclic reaction system, for some values of *S*_*tot*_ and *E*_*tot*_, if the reaction rate constants satisfy the following condition (*SI*, Section 2.1):

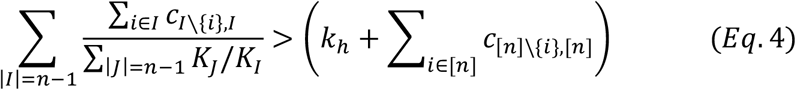

This condition is identical to Eq. 3, but with the extra term *k*_*h*_ on the right-hand side. This is again in line with our analysis above. First, if Eq. 4 holds, then Eq. 3 must also hold, and hence *V*_*S→P*_ is of negative type. Second, Eq. 4 tells us that *k*_*h*_ cannot be larger than a certain amount. This ensures that, with the appropriate choice of *S*_*tot*_ and *E*_*tot*_, *V*_*P→S*_ passes through the last inflection point of *V*_*S→P*_ with slope larger than the slope of *V*_*S→P*_ at that point. As discussed, this gives rise to multiple steady states.

### Conditions for multiple steady states exist for different reaction systems involving a multi-site enzyme

Using the same approach as above, we expanded our analysis to other realistic reaction motifs featuring a multi-site enzyme. We considered two common motifs, involving an enzymatic back reaction from the product to substrate or in- and out-fluxes of both substrate and product (Fig. 3A). The former case represents two enzymes creating a cyclic reaction motif and is commonly found in metabolism and in signalling systems (2,14,15,19). The latter case represents another widely applicable scenario, where any upstream and downstream reactions can generate or consume the substrate and product. In this case, there is no assumption of total substrate amount being conserved.

For each of the cases depicted in Figure 3A, we found that the existence of multiple steady states is guaranteed by an inequality almost identical to Eq. 4 (see *SI* sections 2.2 and 2.3). In the case of a reaction system with an enzymatic back reaction from product to substrate, the catalytic rate constant of the back reaction replaces *k*_*h*_ in Eq. 4. In the case of the reaction system involving fluxes of substrate and product, *k*_*h*_ is eliminated entirely from the inequality, that is, the inequality reduces simply to Eq. 3. These resulting inequalities need to be supplemented with a distinct choice of additional parameters. For the system with enzymatic back reaction, Eq. 4 guarantees multiple steady states after appropriately selecting *S*_*tot*_, *E*_*tot*_ and the conserved total amount of the enzyme catalysing the back reaction from product to substrate (*SI*, Section 2.2). For the system with fluxes, Eq. 3 guarantees multiple steady states after appropriately choosing *E*_*tot*_ and flux rate constants (*SI*, Section 2.3). So, in this case, the possibility of multiple steady states is not conditioned on the value of *S*_*tot*_ as the total amount of substrate is no longer conserved.

The key, intuitive message, as depicted in Fig. 3B, is that a key sufficient mechanism for existence of multiple steady states is related to the dynamics of two distinct sets of enzyme complexes, those that are fully-bound and those that have one binding site empty. When the kinetics of the latter dominates over that of the former, and Eq. 3 is satisfied, a negative type *V*_*S→P*_ curve emerges from the multi-site enzyme dynamics and multiple steady states are guaranteed to exist in some parameter regime in the system.

It is important to note that especially with increasing *n* many multiple steady states may arise, and not just three. We note that a formal analysis of the stability of each steady state cannot be done using the presented general framework. In the case of systems with *n* = 2 and 3, we have sampled kinetic parameter values satisfying Eq. 4, and found that when the system displays three steady states, then bistability arises, showing that at least two steady states are stable. Finally, we also note, that Eq. 3 and 4 do not define *necessary* conditions for multiple steady states, but rather conditions that *guarantees* multiple steady states. As we argued above, there can be parameter sets that lead to a positive type *V*_*S→P*_ curve with multiple peaks and therefore still lead to multiple steady states without fulfilling Eq. 4 (*SI*, Section 2.4).

### Enzyme parameters in the physiological ranges that satisfy Eq. 3 permit bistability

As described above, Eq. 4 describe conditions on the catalytic rates and *K*_*m*_ constants that are guaranteed to result in multiple steady states for *some* set of *S*_*tot*_ and *E*_*tot*_ values. To identify ranges of these latter parameters, we used numerical and analytical methods with the 2-site enzyme model with a cyclic reaction motif involving a non-enzymatic back reaction as a case study (first panel of Fig. 3A). We have chosen kinetic parameters in a physiological range using available information from the literature on multi-site enzymes involved in cyclic reaction systems (see *Methods*). We then derived a bifurcation diagram for the parameters *S*_*tot*_ and *E*_*tot*_ (see *Methods*). We find that for physiologically relevant kinetic parameters, there is a relatively wide range of *S*_*tot*_ and *E*_*tot*_ values allowing for multiple steady states, but *E*_*tot*_ is always much smaller than *S*_*tot*_ (Fig. 4A, red area bounded by dashed lines). In other words, the manifestation of multiple steady states in this cyclic reaction scheme happens in a regime of substrate-saturated enzymes. In fact, for this reaction system, we find that the relation *S*_*tot*_ > *n E*_*tot*_ needs to hold for systems satisfying Eq. 4 to display multiple steady states (see *SI*, Section 2.1).

**Figure 4.**
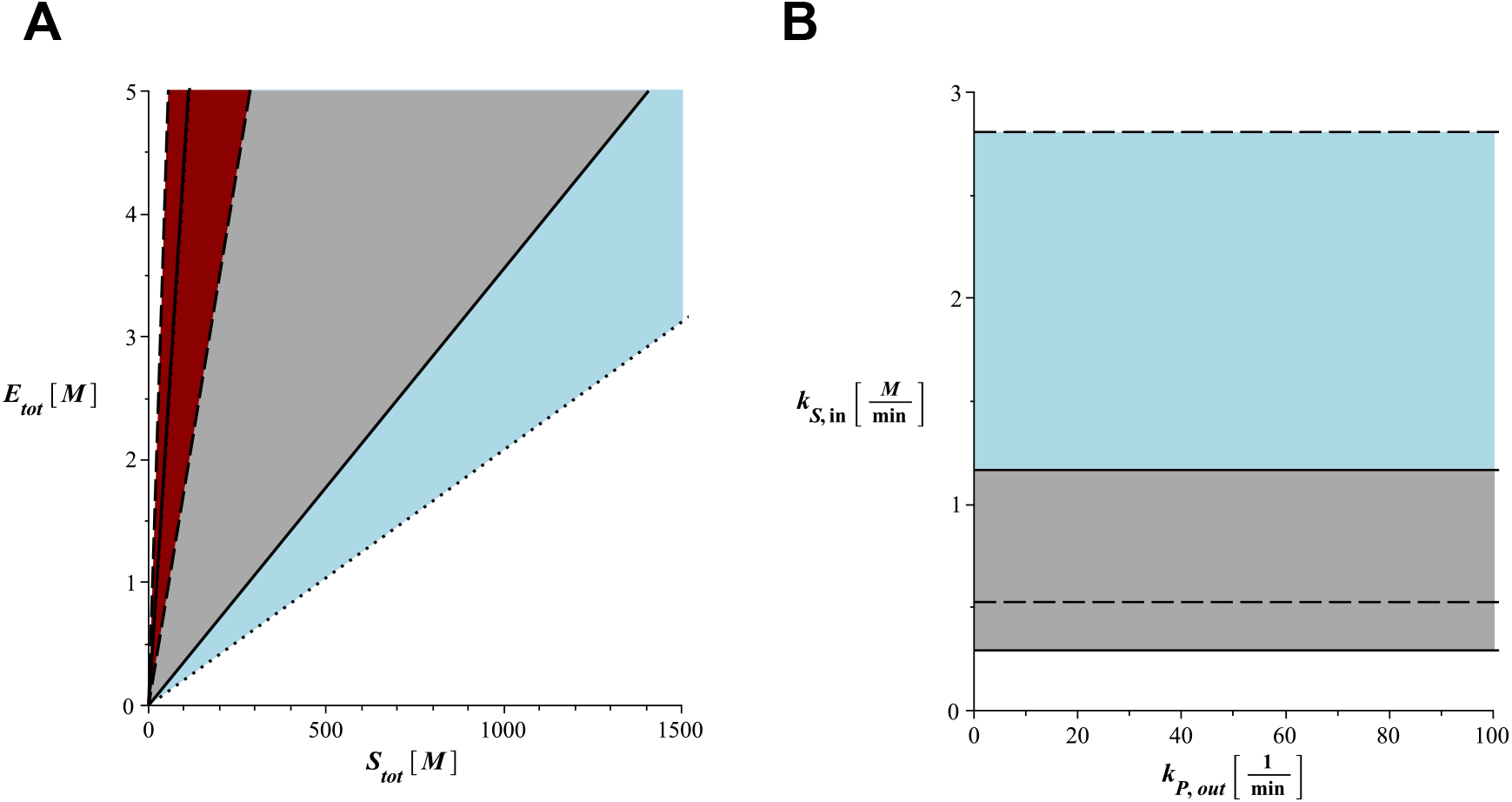
**A**. Two-parameter bifurcation diagram for the reaction system shown on the inset of Fig. 2C and involving a 2-site enzyme with a *P* to *S* back reaction. The diagram shows the regime with three steady states for varying *S*_*tot*_ (x-axis) and *E*_*tot*_ (y-axis) values (in M) for three different sets of physiologically relevant enzyme kinetic parameters. The kinetic parameters used for the region bounded by the dashed lines (covering all of the red area) were: *k*_*1*_ = *k*_*4*_ = *k*_*6*_ = *k*_*10*_ = 10^8^ M^-1^min^-1^, *k*_*2*_ = *k*_*5*_ = *k*_*7*_ = *k*_*11*_ = 10^4^ min^-1^, *k*_*3*_ = 10^5^, *k*_*12*_ = 1.5 10^5^ min^-1^, *k*_*8*_ = *k*_*13*_ = 10^3^ min^-1^, *k*_*h*_ = 0.5·10^3^ min^-1^. For the region bounded by the straight lines (covering all of the grey area and some of the red area), the only parameter altered was the hydrolysis rate of the product; *k*_*h*_ =10^2^ min^-1^. For the region bounded by the dotted lines (covering all the blue and grey areas, and some of the red area), the two parameters altered were the hydrolysis rate of the product and the catalytic rate of one of the single-bound complex; *k*_*h*_ = 10^2^ min^-1^ and *k*_3_ = 10^6^. Note that the left boundary of the regions bounded by the straight and dotted lines overlap. **B**. Two-parameter bifurcation diagram for the reaction system with free substrate and product fluxes (as shown on the right most cartoon on Fig. 3A) and involving a 2-site enzyme. The diagram shows the regime with three steady states for varying substrate in-flux rate *k*_*S,in*_ (y-axis) and product out-flux rate, *k*_*P,out*_ (x-axis). Parameters used were as for the straight-line case of part A, and with additional parameters set as; *k*_*S,out*_ = 10 min^-1^, *k*_*P,in*_ = 0 (no product in-flux). The parameter *E*_*tot*_ was set to 4.15 10^− 5^ M and 10^−4^ M for the areas bounded by the straight and dashed lines respectively.

How would changing kinetic parameters affect the *S*_*tot*_ and *E*_*tot*_ ranges permitting multiple steady states? As discussed above, *S*_*tot*_ determines the intersection point of the *V*_*P→S*_ line with the x-axis, while *E*_*tot*_ determines the height of the *V*_*S→P*_ curve. We can therefore expect that kinetic parameters affecting the slope and shape of the *V*_*P→S*_ line and the *V*_*S→P*_ curve will alter the *S*_*tot*_ and *E*_*tot*_ ranges permitting multiple steady states. In line with this prediction, we find that decreasing *k*_*h*_ and increasing the catalytic rates of the non-fully-bound enzyme complexes widens the *S*_*tot*_ and *E*_*tot*_ range for multiple steady states (Fig. 4A, regions bounded by straight and dotted lines). The latter creates this effect by changing the slope of the *V*_*P→S*_ line, while the latter by changing the height of the *V*_*S→P*_ curve.

In the case of the reaction system with substrate and product fluxes (Fig. 3A, left-most panel), i.e. where *S*_*tot*_ is not a constant anymore, the bistable regime is determined by enzyme kinetic parameters, substrate in- and out-flux, product out-flux, and *E*_*tot*_ (see *SI* section 2.3). For this case, we derived a bifurcation diagram for substrate in-flux and product out-flux rates for a given, physiologically realistic *E*_*tot*_ and found that changing *E*_*tot*_ can result in widening of the bistable regime for these two parameters (Figure 4B).

## METHODS

### Core biochemical model

We considered first a core model involving an enzyme with multiple substrate-binding sites, each able to convert the substrate into a product, as shown in Fig. 1. For this model we assumed that the total enzyme concentration and the total substrate concentration, that is free substrate, substrate bound to enzyme, and the product, are conserved. We relaxed the latter assumption in subsequent models that were built from this core model. For the core model, the resulting binding and catalytic reactions for an enzyme with *n*–binding sites is given in Eq. 1. Additional reactions in the subsequent models and involving the product, and sometimes the substrate, are considered, either as occurring with a constant rate or mediated by an additional enzyme. Our mathematical analyses consisted of writing ordinary differential equations (ODEs) for such reaction systems using mass action kinetics. The ODEs for the core, general model shown in Fig. 1, as well as the alternative models shown in Fig. 3, are provided in full in the *SI* along with the detailed derivations leading to Eq. 2, Eq. 3 and Eq. 4. As an illustration, we provide here the reaction system for the core model, for *n* = 2, i.e. a two-binding-site enzyme:

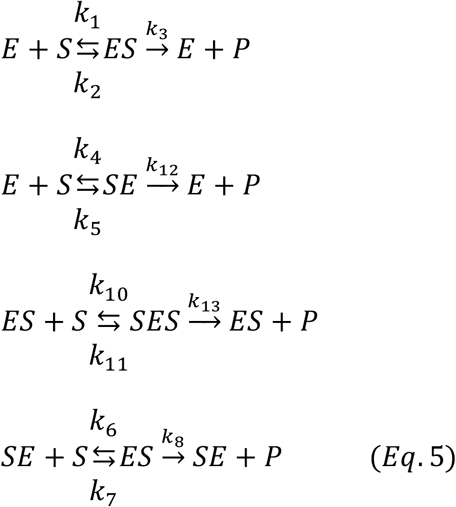

where the single- and double-bound enzyme complexes are denoted as *ES, SE*, and *SES* respectively. The corresponding set of ODEs resulting from this reaction system can be written using mass action kinetics for each of the reactions shown in Eq. 4, as we have done in the provided MATLAB code (see *SI* file1). The conservation relations for this system are:

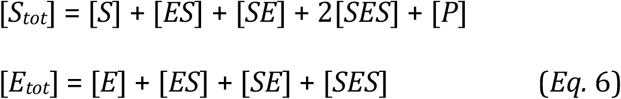

### Symbolic and numerical computations

For all symbolic computations, utilised in finding steady state solutions and deriving mathematical conditions on rate parameters, we used the software Maple 2020. For simulations, run to numerically analyse select systems, we again used Maple, or the MATLAB package, with the standard solver functions.

### Bifurcation analysis and physiologically realistic kinetic parameters and *S*_*tot*_ and *E*_*tot*_ ranges

To analyse if multiple steady states would be realised in physiologically realistic parameter regimes, we used a cyclic reaction system with a two-binding site enzyme (Fig. 4A). For such an enzyme, we have used kinetic parameter values in physiologically feasible ranges as found in the literature and listed below (16,20,21). We then used our mathematical condition shown in Eq. 4, and bifurcation analyses to derive the *S*_*tot*_ and *E*_*tot*_ ranges that guarantee multiple steady states. The analysis was performed using cylindrical algebraic decomposition in Maple, using the package RootFinding[parametric] (22). As an example, the kinetic rate values used for Fig. 2, as listed on its legend, result in Eq. 4 to be satisfied and hence would result in multiple steady states when combined with any combination of *S*_*tot*_ and *E*_*tot*_ that are in the permissible range shown in Fig. 4. The literature based, physiologically realistic kinetic parameter ranges we have considered were: 10^7^ – 10^10^ M^-1^ min^-1^ for substrate-enzyme binding, 10^2^ – 10^6^ min^-1^ for substrate dissociation from a substrate-enzyme complex, 50 – 10^7^ min^-1^ for catalytic rates of enzyme complexes and hydrolysis rate (i.e. *k*_*h*_), and 10^−6^ – 10^−2^ M for their *K*_*M*_values. The literature based, physiologically realistic values of *S*_*tot*_ and *E*_*tot*_ that we considered were 10^−6^ – 10^−2^ M and 10^−8^ – 10^−4^ M respectively.

## DISCUSSION

We have shown that multi-substrate binding enzymes have an inherent capacity to generate bistability when placed within a reaction system. Specifically, the very act of an enzyme binding two or more molecules of the same substrate is guaranteed to result in a specific nonlinear relation between substrate amount and catalytic flux rate (*V*_*S→P*_) in a certain parameter regime (we called the resulting relation a negative type curve in the main text). When the multi-substrate enzyme is placed within a reaction system, this inherent dynamical feature of a negative type curve then guarantees the emergence of multiple steady states. The wider reaction systems, embedding a multi-site enzyme can involve either substrate-product-substrate cycles or systems involving open substrate and product fluxes arising.

These types of reaction systems, as well as multi-site enzymes embedded in them, are common occurrences in metabolic and signalling pathways. Dehydrogenases and kinases, for example, are commonly involved in substrate-to-product cycles (as shown in Fig. 3A), either via redox cycling or phosphorylation/dephosphorylation of substrate-product pairs. Examples include reactions involving dehydrogenases such as lactate or glutamate dehydrogenase (23), and kinase/phosphatase pairs such as those involved in the conversion of fructose-6-phosphate (24). The case with substrate and product fluxes (Fig. 3A, left panel) is a particularly generic scenario, where there is no mass conservation assumption with regards to the substrate and product, and no requirement for a cyclic reaction motif. In these different, common reaction systems, we demonstrate that a multi-site enzyme can lead to bistable dynamics. This is because the negative type *V*_*S→P*_ curve is an inherent feature of the multi-site enzyme and therefore independent of downstream product (and substrate) conversions. Thus, any arrangement of a reaction system resulting in substrate and product conversion dynamics that is capable of intersecting a *V*_*S→P*_ curve of a negative type three times, will result in a system capable of multiple steady states, as we show here.

To directly ascertain bistable parameter regimes, we derived here a mathematical inequality (Eq. 3) that guarantees the *V*_*S→P*_ to be of the negative type. This inequality constitutes the core part of additional inequalities (see Eq. 4 and *SI*) that are derived for different, and common, scenarios of reaction systems embedding a multi-site enzyme, and that guarantee the existence of multiple steady states in them. A key, biochemical intuition arising from these mathematical inequalities is that bistability within a system containing a multi-site enzyme requires non-fully-bound enzyme complexes to ‘outcompete’ the fully-bound complex in terms of catalysis (or flux) from substrate to product. This relates our work to the concept of ‘substrate inhibition’, which is observed in the case of many multi-site enzymes and specifically dehydrogenases and kinases (25), and which is commonly attributed to allosteric effects (i.e. substrate binding also at a non-catalytic, regulatory site on the enzyme). In our case, we emphasize that we do not consider allosteric effects, however, we note that the dynamics we describe here would produce a similar effect as the commonly observed reduction in catalytic rate with increasing substrate concentration (i.e. substrate inhibition). Indeed, when the criteria on kinetic parameters given in Eq. 3 are fulfilled, the resulting dynamics of catalysis rate with increasing substrate concentration (as shown in Fig 2A) will be similar as seen with substrate inhibition. Whether, in the case of specific, natural enzymes displaying substrate inhibition, the fully-bound enzyme complexes have indeed specifically lower catalytic rates than complexes with non-fully-bound complexes, needs to be determined through kinetics experiments.

In addition to the presented inequalities to be satisfied, bistability also requires additional system parameters to be chosen appropriately when we consider systems with cyclic reaction motifs. We find that these additional system parameters, total substrate and enzyme concentrations, as well as kinetic rate constants of additional reactions leading to bistability, exist within physiologically feasible parameter values obtained from enzymatic studies. A key aspect that we note, in the case of cyclic system, is that total substrate levels (i.e. substrate and product combined) need to be larger than total enzyme concentration. This condition is found to be satisfied for many enzymes *in vivo* (21). In line with these findings demonstrating physiological feasibility, bistability in systems involving cyclic reaction motifs are observed when multi-site enzymes are re-constituted *in vitro*, for example using pyruvate kinase, lactate dehydrogenase, or isocitrate dehydrogenase enzymes and their corresponding partners, bistability has been demonstrated experimentally (15,23,26). In the case of systems with open substrate and product fluxes, Eq. 3 guarantees multiple steady states after appropriately choosing *E*_*tot*_ and flux rate constants. Interestingly, in this case, we find that tuning of total enzyme levels, which can be implemented with gene expression control, can widen, or limit the bistable parameter regime. Therefore, our findings of bistability and the parameter regimes it is manifested in, can be of wide relevance for the study of a large range of cellular reaction systems.

Reaction system dynamics are implicated to possess a level of autonomous regulation (1,2). Our findings show that multi-site enzymes can indeed provide such regulation by providing reaction systems with the capability of bistability. When bistability is realised, this will manifest itself as two different steady state concentrations, among which the system can quickly switch. Multi-site enzymes can provide a simple mechanism to achieve such higher-level functions. To this end, our findings provide clear experimental routes towards generating or removing bistability in natural reaction systems or engineered enzymes through the control of kinetic parameters or expression levels with synthetic biology approaches (27). The engineering principles described here for bistability can be further extended to explore the possible sources of multistability and oscillatory dynamics, both of which are observed in models with multi-site enzymes with flux (2,14,28), through further mathematical approaches.

