## Supplementary Information for "Multi-site enzymes as a mechanism for bistability in reaction networks"

#### 1 Steady states of a multi-site enzyme

##### 1.1 Steady state expressions

We consider a multi-site enzyme in which the substrate binds the enzyme at $n$ sites. We let $S$ be the substrate, $E$ the enzyme. We consider the sites identifiable with numbers 1 to $n$, such that we can refer to the first site, second site, and so on. For each subset $I \subseteq \{1,\dots,n\}$, we denote by $ES_I$ the enzyme-substrate complex where the substrate is bound to $E$ at the sites with the numbers in $I$. For example, $ES_{\{1,3\}}$ denotes the complex in which the substrate is bound to the enzyme at the first and third sites; likewise $ES_{\{2\}}$ is the complex in which the substrate is bound to the enzyme only at the second site. The number of elements of $I$ indicates the number of occupied sites. To lighten the notation, we write $[n] = \{1,\dots,n\}$, which is the set for the fully bound enzyme.

The reaction mechanism under consideration comprises the following reactions:

$$S + E \underset{b_{0,i}}{\overset{a_{0,i}}{\rightleftharpoons}} ES_{\{i\}} \xrightarrow{c_{0,i}} P + E, \qquad i = 1,\dots,n \tag{1.1}$$

$$S + ES_I \underset{b_{I,J}}{\overset{a_{I,J}}{\rightleftharpoons}} ES_J \xrightarrow{c_{I,J}} P + ES_I \quad I, J \subseteq [n] \text{ and the set difference between } J \text{ and } I \text{ is one element.}$$

Here $I$ and $J$ are nonempty, that is, each contains at least one index. We identify $E$ with $ES_\emptyset$, the complex with the empty set as subindex.

Throughout this text, we use brackets (e.g. $[E]$, $[S]$) to denote the concentrations of these species.

There are two conservation relations, one for the enzyme and one for the substrate:

$$E_{\text{tot}} = [E] + \sum_{I\subseteq[n], I\neq\emptyset} [ES_I], \tag{1.2}$$

and

$$S_{\text{tot}} = [S] + [P] + \sum_{I\subseteq[n], I\neq\emptyset} |I|\,[ES_I], \tag{1.3}$$

where $|I|$ is the number of elements of $I$, aka the number of occupied sites in $ES_I$. In Section 2.1 and 2.2 of this text, we treat $E_{\text{tot}}$ and $S_{\text{tot}}$ as parameters. We consider the mass-action ODE system defined by the reactions above and the corresponding positive steady states constrained by the two conservation relations.

**Theorem 1.** *Define $M = 2^n - 1 - n$ and consider the system comprising the steady state equations $[\dot{ES_J}] = 0$ for all $J \subseteq [n]$, and the conservation law of enzyme* (1.2)*. Then, the solution to this system is*

$$[ES_I] = \frac{E_{\text{tot}} \sum_{\ell=|I|}^{M+|I|} \alpha_{I,\ell}\, [S]^\ell}{\sum_{J\subseteq[n]} \sum_{\ell=|J|}^{M+|J|} \alpha_{J,\ell}\, [S]^\ell}, \qquad \textit{for all } I \subseteq [n], \tag{1.4}$$

*where all coefficients $\alpha_{I,\ell}$, for any $I \subseteq [n]$ and $\ell$, are nonnegative functions of the kinetic reaction constants for the reactions in* (1.1)*. Furthermore,*

*(1) For $I \subseteq [n]$, $[ES_I]$ is smaller than $E_{\text{tot}}$ for any $[S]$.*

*(2) $[ES_{[n]}]$ goes to $E_{\text{tot}}$ when $[S]$ goes to infinity and increases in $[S]$ for $[S]$ large enough.*

*(3) For $I \subseteq [n]$ with $I \neq [n]$, $[ES_I]$ decreases towards zero as $[S]$ goes to infinity.*

(4) *For $I \subseteq [n]$ with $I \neq \emptyset$, $[ES_I]$ is zero when $[S]$ is zero.*

(5) *If $I$ has $n-1$ elements, then $\alpha_{I,M+n-1} = K_I\, \alpha_{[n],M+n}$, where $K_I = \frac{b_{I,[n]}+c_{I,[n]}}{a_{I,[n]}}$ is the Michaelis-Menten constant of the reaction scheme $S + ES_I \rightleftharpoons ES_{[n]} \longrightarrow P + ES_I$.*

(6) *We let $Ssum := [S] + \sum_{I\subseteq[n], I\neq\emptyset} |I|\, [ES_I]$. Then $Ssum$ increases in $[S]$ for $[S]$ large enough and is zero when $[S]$ is zero.*

*All the statements remain true if all or some of the catalytic kinetic reaction constants $c_{I,J}$ are set to zero, or if all or some of the kinetic reaction constants $b_{I,J}$ are set to zero.*

*Proof.* In what follows, we write $\alpha_{*,*}$, $a_{*,*}$, $b_{*,*}$ and $c_{*,*}$ to generically refer to the respective indexed families of parameters. The set of species formed by $E$ and all enzyme-substrate complexes is a *non-interacting set* in the sense of [1]. This means that no pair of species appears together at any side of a reaction. For any such set, [1] gives explicitly the solution to the steady state equations of the species in this set and the conservation relations among the species in the set in terms of the remaining concentrations. In our setting, the procedure, which we apply now, gives an explicit description of the concentrations of the species at a positive steady state in terms of the concentration of the substrate.

The first step is to construct a digraph $\mathcal{G}$ with vertices $E$ and $ES_I$ for all $I \subseteq [n]$ and the following labeled directed edges:

$$ES_I \xrightarrow{a_{I,J}[S]} ES_J, \qquad ES_J \xrightarrow{b_{I,J}+c_{I,J}} ES_I,$$

and

$$E \xrightarrow{a_{0,i}[S]} ES_{\{i\}}, \qquad ES_{\{i\}} \xrightarrow{b_{0,i}+c_{0,i}} E,$$

for $I, J \subseteq [n]$ such that the set difference $[J] \setminus [I]$ has one element. As shown, if the target complex of an edge has a higher number of occupied sites than the source complex, then the label of the edge is a kinetic reaction constant times $[S]$. If that is not the case, then the label is a reaction rate constant and hence is constant in $[S]$. It is useful to think of the graph as structured into levels depending on the cardinality of $I$: at the 0-th level, we have $E$; at the 1st level, we have all complexes where $I$ has one element, and so on. The $n$-th level is $ES_{[n]}$. See Figure S1 for an example with $n = 3$. The label of all edges pointing downwards, increasing the level, have $[S]$ as a multiplier, while the labels of the edges pointing upwards, decreasing the level, do not involve $[S]$.

We consider spanning trees rooted at a vertex $V$: this is a tree that contains all vertices of the graph $\mathcal{G}$, and such that there is a directed path from each vertex to $V$. Then the vertex $V$ is the only

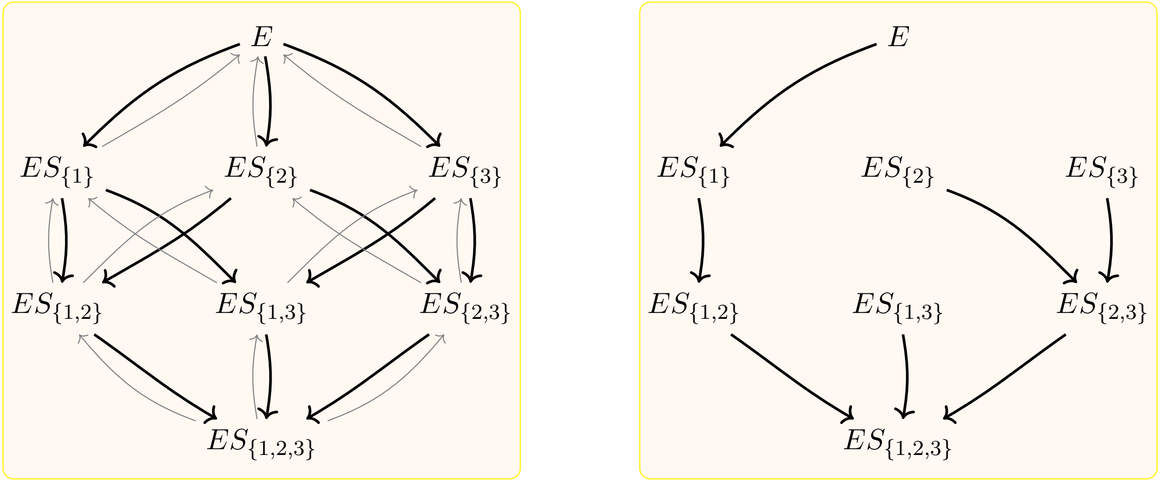


Figure S1: Left panel: Example of the graph $\mathcal{G}$ in the proof of Theorem 1 for $n = 3$. Thick edges have labels of the form $a_{*,*}[S]$, while light edges have labels of the form $b_{*,*} + c_{*,*}$. Labels are omitted for clarity. Right panel: Example showing one of the spanning trees rooted at $ES_{\{1,2,3\}}$.

vertex with no outgoing edge. All other edges have exactly one outgoing edge. For each such tree $T$, we denote by $\pi(T)$ the product of the labels of the edges in the tree. We define $A$ to be the sum of $\pi(T)$ for all possible spanning trees of $\mathcal{G}$ rooted at any vertex, and $A(V)$ to be the sum of $\pi(T)$ for all spanning trees rooted at $V$. In particular, $A = \sum_V A(V)$.

By [1], at steady state it holds

$$[ES_I] = \frac{E_{\text{tot}} A(ES_I)}{A}. \tag{1.5}$$

Since the labels are either a positive constant or a positive constant times the concentration of $[S]$, this becomes a rational function in $[S]$ with all non-zero coefficients positive. Therefore, to study $[ES_I]$, we need to understand the spanning trees of $\mathcal{G}$. In particular, we study the maximal and minimal number of edges with $[S]$ that any such tree can have, which determines the degree of $A(V)$ and the smallest exponent of the monomials of $A(V)$.

We start with some observations:

- $2^n$ is the possible number of subsets of $[n]$, which corresponds to the number of vertices of $\mathcal{G}$. Each spanning tree has exactly $2^n - 1$ edges, and hence the degree of $A(V)$ in $[S]$ can at most be $2^n - 1$.

- All edges outgoing $E$ have a label which is a multiple of $[S]$. This implies that for all vertices $V \neq E$ of the form $ES_I$, any tree rooted at $V$ has at least one edge with label a multiple of $[S]$. As a consequence, $A(V)$ has no independent term in $[S]$ and is zero when $[S] = 0$ (showing statement (4)).

We start by looking at the case $I = [n]$, such that $V = ES_{[n]}$. In this case, for any vertex other than $V$, choose an edge with label multiple of $[S]$, that is, increasing the level. Any such choice gives rise to a spanning tree $T$ rooted at $V$ and $\pi(T)$ is a multiple of $[S]^{2^n-1}$. Thus $A(V)$ is a polynomial of degree $2^n - 1$ in $[S]$. Note that $2^n - 1 = M + n$, which is the degree of the numerator of (1.4) for $ES_{[n]}$.

Now consider any other set $I \subseteq [n]$. Pick a spanning tree rooted at $ES_{[n]}$ as explained above. Then there is a unique path from $ES_I$ to $ES_{[n]}$, which consists only of edges increasing the level and hence all labels are multiplied by $[S]$. This path has $n - |I|$ edges, because each edge increases one level. If this particular path is reversed by replacing each edge in the path with its reverse edge, then we obtain a spanning tree rotted at $ES_I$ rather than $ES_{[n]}$. For this new tree, the reversed path has labels of the form $b_{*,*} + c_{*,*}$ rather than involving $[S]$. Thus, the exponent of $[S]$ in the product of the labels of the new tree rooted at $ES_I$ equals the exponent of the tree rooted at $ES_{[n]}$ minus the number of edges in the reversed path. This gives that the label of this spanning tree is a multiple of $[S]^{2^n-1-n+|I|} = [S]^{M+|I|}$. Hence $A(V)$ has degree at least $M + |I|$. Any other spanning tree has a path from $ES_{[n]}$ to $ES_I$. This path must have at least $n - |I|$ edges with label $b_{*,*} + c_{*,*}$, because it needs to decrease $n - |I|$ levels. It could have more, if it decreased too much and then increased. This means that the degree of $A(V)$ cannot be larger than $M + |I| = (2^n - 1) - (n - |I|)$, since there cannot be more than $M + |I|$ edges whose label is multiple of $[S]$. This shows that the degree of the numerator of (1.5) is as given in (1.4)

We now find the smallest of the exponents of the monomials in the numerator of (1.5). We argue similarly, but focusing on the vertex $E$ instead of $ES_{[n]}$. For any vertex other than $E$, choose an edge decreasing the level, and hence with label not involving $[S]$. These edges define a spanning tree $T$ rooted at $E$, with $\pi(T)$ not depending on $[S]$. Hence, $A(E)$ is a polynomial with smallest exponent 0, which is the cardinality of the empty set. Now consider any vertex $ES_I$, and reverse the edges of the path from $ES_I$ to $E$ in this spanning tree we just constructed. This path has length $|I|$. The new tree is rooted at $ES_I$, and the label is a multiple of $[S]^{|I|}$. Since any path from $E$ to $ES_I$ in a spanning tree rooted at $ES_I$ must involve at least $|I|$ edges with label involving $[S]$, this shows that the smallest exponent in $A(ES_I)$ is $|I|$, as in (1.4). This concludes the proof of the form of $[ES_I]$ as a function of $[S]$, that is, of equation (1.4).

It is clear that $[ES_I]$ goes to zero when $[S]$ goes to infinity and $I \neq [n]$, because the degree of the numerator is smaller than the degree of the denominator, and hence statement (3) holds. For $I = [n]$, the numerator and the denominator of (1.4) have the same degree, and the leading coefficient of the numerator of (1.4) agrees with the leading coefficient of the denominator of (1.4) times $E_{\text{tot}}$. This implies that the limit is $E_{\text{tot}}$ in this case, showing part of statement (2). Since the numerator of $[ES_I]$ in (1.4) divided by $E_{\text{tot}}$ is smaller than the denominator, $[ES_I]$ is smaller than $E_{\text{tot}}$ for all values of $[S]$, and statement (1) holds. Using this, for $[S]$ large enough, $[ES_{[n]}]$ must increase, completing the proof of statement (2).

To show statement (5), note that $\alpha_{I,M+n-1}$ is the sum of the labels of the trees obtained from a spanning tree rooted at $ES_{[n]}$ as explained above, with all edges involving $[S]$, removing the only edge from $ES_I$ to $ES_{[n]}$, which has label $a_{I,[n]}[S]$, and adding the only edge from $ES_{[n]}$ to $ES_I$, which has label $b_{I,[n]} + c_{I,[n]}$. Hence, $\alpha_{I,M+n-1} = \frac{b_{I,[n]}+c_{I,[n]}}{a_{I,[n]}} \cdot \alpha_{[n],M+n} = K_I \cdot \alpha_{[n],M+n}$.

Finally, to see statement (6), from statement (4) we readily see that $Ssum$ is zero when $[S]$ is zero. As $[S]$ goes to infinity, $[ES_{[n]}]$ goes to $E_{\text{tot}}$ and $[ES_I]$ to zero. Hence, $Ssum$, as a function of $[S]$, increases for $[S]$ large enough and goes to infinity. □

**Alternative binding orders.** If some reactions are removed such that there is still at least one spanning tree rooted at $E$ and one rooted at $ES_{[n]}$ in the graph $\mathcal{G}$ in the proof of Theorem 1, then the expressions in Theorem 1 remain true for the enzyme-substrate complexes this smaller network has. For instance, consider ordered binding such that the only complexes that are formed are those with index set $I$ equal to $\{1\}, \{1,2\}, \{1,2,3\}, \dots, [n]$. The reaction scheme is then

$$S + E \rightleftharpoons ES_{\{1\}} \longrightarrow P + E, \quad S + ES_{\{1,\dots,i\}} \rightleftharpoons ES_{\{1,\dots,i+1\}} \longrightarrow P + ES_{\{1,\dots,i\}}, \; i = 1, \dots, n-1.$$

Only the complexes $ES_{\{1,\dots,i\}}$ for $i = 1, \dots, n$ are formed. The graph $\mathcal{G}$ in the proof of Theorem 1 is

$$E \rightleftharpoons ES_{\{1\}} \rightleftharpoons \dots \rightleftharpoons ES_{\{1,\dots,n-1\}} \rightleftharpoons ES_{\{1,\dots,n\}},$$

which has a spanning tree rooted at $E$ and one rooted at $ES_{\{1,\dots,n\}}$. Hence, at steady state, the expressions in Theorem 1 for $[ES_{\{1,\dots,i\}}]$, for $i = 1, \dots, n$, as well as the statements about them, remain valid, by letting $M$ be the number of complexes minus $n + 1$.

**Case $n = 2$.** For $n = 2$, we can show that $[ES_{\{1,2\}}]$ is an increasing function of $[S]$. In this case we have

$$[ES_{\{1,2\}}] = \frac{E_{\text{tot}}\ (\alpha_{\{1,2\},2}[S]^2 + \alpha_{\{1,2\},3}[S]^3)}{\alpha_{\{0\},0} + \alpha_{\{0\},1}[S] + \alpha_{\{1\},1}[S] + \alpha_{\{1\},2}[S]^2 + \alpha_{\{2\},1}[S] + \alpha_{\{2\},2}[S]^2 + \alpha_{\{1,2\},2}[S]^2 + \alpha_{\{1,2\},3}[S]^3}.$$

The derivative of this function (computed in `Maple`) is a quotient of polynomials in $[S]$, such that both the numerator and denominator are positive for positive $[S]$. Hence the function $ES_{\{1,2\}}$ is increasing.

For $[ES_{\{1\}}]$, we have

$$[ES_{\{1\}}] = \frac{E_{\text{tot}}\ (\alpha_{\{1\},1}[S] + \alpha_{\{1\},2}[S]^2)}{\alpha_{\{0\},0} + \alpha_{\{0\},1}[S] + \alpha_{\{1\},1}[S] + \alpha_{\{1\},2}[S]^2 + \alpha_{\{2\},1}[S] + \alpha_{\{2\},2}[S]^2 + \alpha_{\{1,2\},2}[S]^2 + \alpha_{\{1,2\},3}[S]^3}.$$

In this case the derivative with respect to $[S]$ is a quotient of polynomials in $[S]$, the denominator is positive, and the numerator is a polynomial in $[S]$ with one sign change. By the Descartes' rule of signs, it has exactly one root, where the function has a maximum. The concentration of the complex $[ES_{\{2\}}]$ is analysed analogously.

By a similar computation, we express Ssum $= [ES_{\{1\}}] + [ES_{\{2\}}] + [ES_{\{1,2\}}] + [S]$ as a function of $[S]$, compute the derivate in `Maple`, and find that it is a quotient of polynomials with all coefficients positive. Hence, Ssum is an increasing function of $[S]$ for $[S] \geq 0$.

In the main text, we write $ES_{\{1\}}$, $ES_{\{2\}}$ and $ES_{\{1,2\}}$ as SE, ES, SES respectively for simplicity.

#### 1.2 Negative type curves

Recall from the main text that a function with horizontal asymptote at $L$ is said to be of *negative type* if it approaches its limit value from above. We characterize now when $V_{S\to P}$ is of negative type. By definition

$$V_{S\to P} = \sum_{I\subseteq[n], I\neq\emptyset} \sum_{i\in I} c_{I\setminus\{i\},I}[ES_I].$$

By Theorem 1 (2)-(3), $V_{S\to P}$ goes to

$$L := \sum_{i=1}^{n} c_{[n]\setminus\{i\},[n]} E_{\rm tot} \tag{1.6}$$

when $[S]$ goes to infinity.

**Theorem 2.** *Consider the function $V_{S\to P}$ as a function in $[S]$ by using the expressions in Theorem 1. Then $V_{S\to P}$ is of negative type if and only if*

$$\sum_{|I|=n-1} \frac{\sum_{i\in I} c_{I\setminus\{i\},I}}{\sum_{|J|=n-1} \frac{K_J}{K_I}} > \sum_{i\in[n]} c_{[n]\setminus\{i\},[n]}. \tag{1.7}$$

*Proof.* For simplicity, we let $\beta_I = \left(\sum_{i\in I} c_{I\setminus\{i\},I}\right)$ and write $V_{S\to P}$ as $v([S])$. Using the expressions in Theorem 1, we obtain

$$v([S]) = \sum_{I\subseteq[n], I\neq\emptyset} \beta_I [ES_I] = E_{\rm tot} \sum_{I\subseteq[n], I\neq\emptyset} \frac{\beta_I \sum_{\ell=|I|}^{M+|I|} \alpha_{I,\ell}[S]^\ell}{\sum_{J\subseteq[n]} \sum_{\ell=|J|}^{M+|J|} \alpha_{J,\ell}[S]^\ell}.$$

The curve $v([S])$ is of negative type if and only if $v([S])-L$ is positive when $[S]$ goes to infinity. Using that $L=\beta_{[n]}E_{\rm tot}$, we have, after dividing by $E_{\rm tot}$ that

$$\begin{aligned}\tfrac{1}{E_{\rm tot}}(v([S])-L) &= \left(\sum_{I\subseteq[n], I\neq\emptyset} \frac{\beta_I \sum_{\ell=|I|}^{M+|I|} \alpha_{I,\ell}[S]^\ell}{\sum_{J\subseteq[n]} \sum_{\ell=|J|}^{M+|J|} \alpha_{J,\ell}[S]^\ell}\right) - \beta_{[n]} \\ &= \frac{\left(\sum_{I\subseteq[n], I\neq\emptyset} \beta_I \sum_{\ell=|I|}^{M+|I|} \alpha_{I,\ell}[S]^\ell\right) - \beta_{[n]}\left(\sum_{J\subseteq[n]} \sum_{\ell=|J|}^{M+|J|} \alpha_{J,\ell}[S]^\ell\right)}{\sum_{J\subseteq[n]} \sum_{\ell=|J|}^{M+|J|} \alpha_{J,\ell}[S]^\ell}.\end{aligned}$$

The denominator is always positive. This function is thus positive for $[S]$ large enough if and only if the leading coefficient of the numerator is positive. The coefficient of $[S]^{M+n}$ is $\beta_{[n]}\alpha_{[n],M+n} - \beta_{[n]}\alpha_{[n],M+n} = 0$. We compute now the coefficient of $[S]^{M+n-1}$, which comes from the intermediates with $n-1$ bound sites. Using property (5) in Theorem 1, this coefficient is

$$\begin{aligned}\sum_{|I|=n-1} \beta_I \alpha_{I,M+n-1} - \beta_{[n]} \sum_{|I|=n-1} \alpha_{I,M+n-1} &= \sum_{|I|=n-1} (\beta_I - \beta_{[n]})\alpha_{I,M+n-1} \\ &= \sum_{|I|=n-1} \left(\sum_{i\in I} c_{I\setminus\{i\},I} - \sum_{i\in[n]} c_{[n]\setminus\{i\},[n]}\right) K_I\, \alpha_{[n],M+n}. \end{aligned} \tag{1.8}$$

Note that $\sum_{i\in[n]} c_{[n]\setminus\{i\},[n]}$ is the sum of the kinetic constants of the catalytic reactions from $ES_{[n]}$, while $\sum_{i\in I} c_{I\setminus\{i\},I}$ is the sum of the kinetic constants of the catalytic reactions from $ES_I$, that is, the complexes with $n-1$ binding sites.

To wrap up, the term in (1.8) is the leading coefficient of the numerator of $\frac{(v([S])-L)}{E_{\rm tot}}$. When this coefficient is positive, then $v([S])-L$ is positive for all $[S]$ large enough and $v([S])$ approaches $L$ from above. If the coefficient is negative, then the difference is negative for all $[S]$ large enough. Hence, the coefficient being positive is equivalent to $v([S])$) being of negative type. We factor out $\alpha_{[n],M+n}$, which is positive, and obtain the following characterization of negative type curves, written in different ways:

$$\sum_{|I|=n-1}\left(\sum_{i\in I} c_{I\setminus\{i\},I}\right)K_I > \left(\sum_{i\in[n]} c_{[n]\setminus\{i\},[n]}\right)\left(\sum_{|I|=n-1} K_I\right)$$

or

$$\frac{\sum_{|I|=n-1}\left(\sum_{i\in I} c_{I\setminus\{i\},I}\right)K_I}{\sum_{|I|=n-1} K_I} > \sum_{i\in[n]} c_{[n]\setminus\{i\},[n]}$$

or

$$\sum_{|I|=n-1} \frac{\sum_{i\in I} c_{I\setminus\{i\},I}}{\sum_{|J|=n-1}\frac{K_J}{K_I}} > \sum_{i\in[n]} c_{[n]\setminus\{i\},[n]}.$$

Note that the last inequality is as given in the statement of the theorem. This concludes the proof. □

### 2 Multistationarity in the multi-site system in the presence of additional reaction mechanisms

Theorem 2 gives a characterization of when $V_{S\to P}$ is of negative type, which depends on the kinetic rate constants of the core mechanism. In this section, we show that the negative type shape is enough to give rise to multiple positive steady states, when we consider additional and different reactions involving $S$ and $P$, that expand the system considered above. As we add such reactions, we ensure that they leave the conservation relation (1.2), the rate equations for $[E]$, and the intermediate complexes intact, thereby guaranteeing that Theorem 1 and Theorem 2 remain true.

In all studied cases, the inequality

$$\frac{\sum_{|I|=n-1}\sum_{i\in I} c_{I\setminus\{i\},I}K_I}{\sum_{|I|=n-1} K_I} > \sum_{i\in[n]} c_{[n]\setminus\{i\},[n]}$$

or equivalently

$$\sum_{|I|=n-1} \frac{\sum_{i\in I} c_{I\setminus\{i\},I}}{\sum_{|J|=n-1}\frac{K_J}{K_I}} > \sum_{i\in[n]} c_{[n]\setminus\{i\},[n]}$$

characterizing negative type curves is sufficient for the existence of at least two or three positive steady states, after appropriately choosing the value of the total amounts and remaining kinetic parameters.

#### 2.1 Reverse catalysis from $P$ to $S$ (non-enzymatic)

We start by considering reverse catalysis from $P$ to $S$ (e.g via hydrolysis, de-phosphorylation, or other biochemically plausible reactions). In this case, only one additional reaction $P \to S$ is added to the system, which can be enzyme catalysed or not. We consider the non-enzymatic scenario first (in this section) and the enzymatic case next (in Section 2.2)

We start by considering that a reverse catalysis reaction $P \xrightarrow{k_h} S$ is added to the original system (1.1), such that $V_{P\to S} = k_h[P]$. Then, the conservation relation (1.3) holds. In the following theorem we provide an inequality on the kinetic parameters that ensure that the total amounts of substrate and enzyme can be appropriately chosen such that the system has at least three positive steady states.

**Theorem 3.** *Assume all reaction rate constants are fixed and satisfy*

$$\sum_{|I|=n-1} \frac{\sum_{i\in I} c_{I\setminus\{i\},I}}{\sum_{|J|=n-1}\frac{K_J}{K_I}} > \sum_{i\in[n]} c_{[n]\setminus\{i\},[n]} + k_h. \tag{2.1}$$

*Then, there exist $S_{\text{tot}}$ and $E_{\text{tot}}$ such that the system has at least three positive steady states.*

*Proof.* Considering the expressions in Theorem 1, in order to completely solve for the steady states, we need to consider the differential equation for $[P]$ and the conservation law with $S_{\text{tot}}$. Using the conservation law, we obtain:

$$[P] = S_{\text{tot}} - [S] - \sum_{I\subseteq[n], I\neq\emptyset} |I|[ES_I]. \tag{2.2}$$

Using the differential equation for $[P]$, we have

$$\begin{aligned} 0 &= \sum_{I\subseteq[n], I\neq\emptyset} \sum_{i\in I} c_{I\setminus\{i\},I}[ES_I] - k_h[P] \qquad (2.3)\\ &= \sum_{I\subseteq[n], I\neq\emptyset} \sum_{i\in I} c_{I\setminus\{i\},I}[ES_I] - k_h\left(S_{\text{tot}} - [S] - \sum_{I\subseteq[n], I\neq\emptyset} |I|[ES_I]\right)\\ &= \sum_{I\subseteq[n], I\neq\emptyset} \left(\sum_{i\in I} c_{I\setminus\{i\},I} + k_h|I|\right)[ES_I] - k_h(S_{\text{tot}} - [S]). \end{aligned}$$

Hence, the values of $[S]$ at steady state are the positive solutions to

$$\sum_{I\subseteq[n], I\neq\emptyset} \left(\sum_{i\in I} c_{I\setminus\{i\},I} + k_h|I|\right)[ES_I] = k_h(S_{\text{tot}} - [S]), \tag{2.4}$$

and for each solution, a positive steady state of the network is obtained by using the expressions in Theorem 1 and (2.2). So we aim at showing that (2.4) admits at least three positive solutions when (2.1) holds, and after appropriately choosing $S_{\text{tot}}$ and $E_{\text{tot}}$.

For simplicity and as in the proof of Theorem 2, we let $\beta_I = \left(\sum_{i\in I} c_{I\setminus\{i\},I} + k_h|I|\right)$. We let $v([S])$ be the function obtained from the left side of (2.4) after using the expressions in Theorem 1:

$$v([S]) = \sum_{I\subseteq[n], I\neq\emptyset} \beta_I[ES_I] = E_{\text{tot}} \sum_{I\subseteq[n], I\neq\emptyset} \frac{\beta_I \sum_{\ell=|I|}^{M+|I|} \alpha_{I,\ell}[S]^\ell}{\sum_{J\subseteq[n]} \sum_{\ell=|J|}^{M+|J|} \alpha_{J,\ell}[S]^\ell}.$$

This function is identical to the one studied in Theorem 2, but the two differ in the exact value of $\beta_I$. The limit value of $v([S])$ when $[S]$ goes to infinity is $L := \beta_{[n]}E_{\text{tot}}$, and proceed as in the proof of Theorem 2, see (1.8), $v([S])$ is of negative type if and only if

$$\sum_{|I|=n-1} (\beta_I - \beta_{[n]})\alpha_{I,M+n-1} > 0,$$

which is equivalent to

$$\sum_{|I|=n-1} \left(\sum_{i\in I} c_{I\setminus\{i\},I} + k_h(n-1) - \sum_{i\in[n]} c_{[n]\setminus\{i\},[n]} - k_h n\right) K_I\, \alpha_{[n],M+n} > 0$$

$$\text{that is,} \quad \sum_{|I|=n-1} \left(\sum_{i\in I} c_{I\setminus\{i\},I} - \sum_{i\in[n]} c_{[n]\setminus\{i\},[n]} - k_h\right) K_I > 0. \tag{2.5}$$

Rearranging inequality (2.5), we obtain inequality (2.1). Hence, inequality (2.1) in the statement of the theorem is equivalent to $v([S])$ being of negative type. This gives that the function $v([S])$ satisfies $v(0) = 0$, increases when $[S]$ is small, and for $[S]$ large enough it decreases towards a horizontal asymptote at $L$. This implies that $v([S])$ attains a maximum value above $L$, and has an inflection point where $v([S])$ attains a value larger than $L$.

Now we will argue that $E_{\text{tot}}$ and $S_{\text{tot}}$ can be chosen such that the line $k_h(S_{\text{tot}} - [S])$ goes through the inflection point with slope smaller than the tangent line of $v([S])$ at the inflection point. With this, close to the inflection point but at the left side we have $v([S]) - k_h(S_{\text{tot}} - [S]) > 0$, while at the right side it holds $v([S]) - k_h(S_{\text{tot}} - [S]) < 0$. This implies that (2.4) has three solutions as the line $k_h(S_{\text{tot}} - [S])$ necessarily intersects $v([S])$ at at least three values of $[S]$: one at the inflection point, another one after the inflection point, since when $[S]$ goes to infinity $v([S]) - k_h(S_{\text{tot}} - [S]) > 0$ meaning that the function $v([S]) - k_h(S_{\text{tot}} - [S])$ changes sign after the inflection point and hence must be zero somewhere, and similarly, one before the inflection point, since at $[S] = 0$ we have $k_h S_{\text{tot}} > v(0) = 0$ and again $v([S]) - k_h(S_{\text{tot}} - [S])$ changes sign between zero and the inflection point.

First, note that by increasing or decreasing $E_{\text{tot}}$, $v([S])$ preserves its shape. Let $-T < 0$ be the slope of the tangent line of $v([S])$ at the inflection point. By increasing $E_{\text{tot}}$ if necessary, $T$ can be made larger than $k_h$, since multiplying a function with a number implies that the first derivative also gets multiplied by this number. After doing this, consider the line through the inflection point with slope $-k_h$. By letting $S_{\text{tot}}$ be such that the intersection of this line with the vertical axis is $k_h S_{\text{tot}}$, we have that $k_h(S_{\text{tot}} - [S])$ goes through the inflection point with slope smaller than the tangent line of $v([S])$ at the inflection point. By the arguments above, this concludes the proof. □

**Remark 1.** Note in particular that it follows from the proof of Theorem 3 that $S_{\text{tot}}$ is larger than $L$ (from the proof), that is

$$S_{\text{tot}} > \frac{E_{\text{tot}}}{k_h}\left(\sum_{i\in[n]} c_{[n]\setminus\{i\},[n]} + nk_h\right) = E_{\text{tot}}\left(\frac{1}{k_h}\sum_{i\in[n]} c_{[n]\setminus\{i\},[n]} + n\right),$$

for there to be multiple intersection points between $v([S])$ and $k_h(S_{\text{tot}} - S)$. From this we derive that necessarily

$$S_{\text{tot}} > nE_{\text{tot}}.$$

**Remark 2.** If $V_{S\to P}$ is of negative type, then (1.7) holds, and we can choose $k_h$ small enough such that (2.1) holds. Hence by Theorem 3, the system admits three positive steady states after appropriately choosing $k_h$, $E_{\text{tot}}$ and $S_{\text{tot}}$. Additionally, if (2.1) holds, then necessarily (2.1) also holds and $V_{S\to P}$ is of negative type. Therefore, Theorem 3 gives a condition to find suitable values for $k_h$ provided $V_{S\to P}$ is of negative type.

#### 2.2 Reverse catalysis from $P$ to $S$ (enzymatic)

We assume now that the conversion of $P$ back to $S$ proceeds via an enzymatic mechanism where the new enzyme is denoted by $K$:

$$P + K \underset{k_2}{\overset{k_1}{\rightleftharpoons}} PK \xrightarrow{k_3} S + K.$$

Then, $[PK]$ becomes an extra term in the conservation relation with $S_{\text{tot}}$, (1.3), and there is a new conservation relation $K_{\text{tot}} = [K] + [PK]$.

**Theorem 4.** *Assume that all reaction rate constants are fixed, and that*

$$\sum_{|I|=n-1} \frac{\sum_{i\in I} c_{I\setminus\{i\},I}}{\sum_{|J|=n-1}\frac{K_J}{K_I}} > \sum_{i\in[n]} c_{[n]\setminus\{i\},[n]} + k_3. \tag{2.6}$$

*Then $S_{\text{tot}}, E_{\text{tot}}$ and $K_{\text{tot}}$ can be chosen such that the system has at least three positive steady states.*

*Proof.* The proof follows closely the proof of Theorem 3. At steady state we have

$$[PK] = \alpha[P][K], \qquad \text{where} \quad \alpha = \tfrac{k_1}{k_2+k_3}$$

is the inverse of the Michaelis-Menten constant. This equation together with the conservation relation with $K_{\text{tot}}$, determines $[K]$ and $[PK]$ as functions of $[P]$:

$$[K] = \tfrac{K_{\text{tot}}}{1+\alpha[P]}, \qquad [PK] = \tfrac{\alpha K_{\text{tot}}[P]}{1+\alpha[P]}.$$

Assuming these relations, the differential equation for $[P]$, analogous to (2.3) is

$$[\dot{P}] = \sum_{I\subseteq[n], I\neq\emptyset} \sum_{i\in I} c_{I\setminus\{i\},I}[ES_I] - \tfrac{k_3\alpha K_{\text{tot}}[P]}{1+\alpha[P]}$$

So, at steady state

$$\sum_{I\subseteq[n], I\neq\emptyset} \sum_{i\in I} c_{I\setminus\{i\},I}[ES_I] = \tfrac{k_3\alpha K_{\text{tot}}[P]}{1+\alpha[P]}. \tag{2.7}$$

Furthermore,

$$S_{\text{tot}} = [S] + [P] + \sum_{I\subseteq[n], I\neq\emptyset} |I|[ES_I] + \tfrac{\alpha K_{\text{tot}}[P]}{1+\alpha[P]}. \tag{2.8}$$

We let

$$v_1([S]) = \sum_{I\subseteq[n], I\neq\emptyset} \sum_{i\in I} c_{I\setminus\{i\},I}[ES_I] \quad \text{and} \quad v_2([S]) = \sum_{I\subseteq[n], I\neq\emptyset} |I|[ES_I].$$

After isolating $[P]$ from equation (2.7), we obtain

$$[P] = \frac{v_1([S])}{\alpha(k_3K_{\text{tot}} - v_1([S]))},$$

which is positive provided $k_3K_{\text{tot}} > v_1([S])$. Inserting this value into (2.8), we obtain

$$\frac{v_1([S])}{\alpha(k_3K_{\text{tot}} - v_1([S]))} + v_2([S]) + \frac{v_1([S])}{k_3} = S_{\text{tot}} - [S].$$

The positive steady states are in one to one correspondence with the positive solutions to this equation. We denote by $v([S])$ the left side of this equation and proceed similarly to Section 2.1. We let $L' := \left(\sum_{i\in[n]} c_{[n]\setminus\{i\},[n]}\right)$ be such that the limit of $v_1([S])$ as $[S]$ goes to infinity is $L'\,E_{\text{tot}}$, and then, the limit of $v([S])$ as $[S]$ goes to infinity is

$$L := \frac{L'\,E_{\text{tot}}}{\alpha(k_3K_{\text{tot}} - L'\,E_{\text{tot}})} + nE_{\text{tot}} + \frac{L'\,E_{\text{tot}}}{k_3}.$$

If $k_3K_{\text{tot}} > L'\,E_{\text{tot}}$, then $L$ is guaranteed to be positive.

We study under what conditions $v([S]) - L$ is positive for $[S]$ going to infinity. We have

$$v([S]) - L = \frac{v_1([S])}{\alpha(k_3K_{\text{tot}} - v_1([S]))} - \frac{L'\,E_{\text{tot}}}{\alpha(k_3K_{\text{tot}} - L'\,E_{\text{tot}})} + v_2([S]) - nE_{\text{tot}} + \tfrac{1}{k_3}(v_1([S]) - L'E_{\text{tot}}).$$

Let us first study $B := v_2([S]) - nE_{\text{tot}} + \frac{1}{k_3}(v_1([S]) - L'E_{\text{tot}})$. We use the same argument as in Section 2.1, but now with $\beta_I = \left(\sum_{i\in I} c_{I\setminus\{i\},I} + k_3|I|\right)$, to show that $B$ is positive for $[S]$ large enough if and only if inequality (2.6) in Theorem 4 holds. For the other summand, we have

$$\begin{aligned}\frac{v_1([S])}{\alpha(k_3K_{\text{tot}} - v_1([S]))} - \frac{L'\,E_{\text{tot}}}{\alpha(k_3K_{\text{tot}} - L'\,E_{\text{tot}})} &= \frac{v_1([S])(k_3K_{\text{tot}} - L'\,E_{\text{tot}}) - L'\,E_{\text{tot}}(k_3K_{\text{tot}} - v_1([S]))}{\alpha(k_3K_{\text{tot}} - v_1([S]))(k_3K_{\text{tot}} - L'\,E_{\text{tot}})}\\ &= \frac{k_3K_{\text{tot}}(v_1([S]) - L'\,E_{\text{tot}})}{\alpha(k_3K_{\text{tot}} - v_1([S]))(k_3K_{\text{tot}} - L'\,E_{\text{tot}})}.\end{aligned}$$

Provided the denominator is positive, as $v_1([S]) = V_{S\to P}([S])$ as in Theorem 2, we obtain that $v_1([S]) - L'\,E_{\text{tot}}$ is positive for $[S]$ large enough if and only if

$$\sum_{|I|=n-1} \frac{\sum_{i\in I} c_{I\setminus\{i\},I}}{\sum_{|J|=n-1} \frac{K_J}{K_I}} > \sum_{i\in[n]} c_{[n]\setminus\{i\},[n]} \tag{2.9}$$

holds. But note that if (2.6) holds, then so does (2.9).

So, assume (2.6) holds, such that $v([S]) - L$ is positive for $[S]$ large enough, provided both $k_3 K_{\text{tot}} > L' E_{\text{tot}}$ and $k_3 K_{\text{tot}} > v_1([S])$ hold for all $[S]$. For these two inequalities to hold, it is enough to choose $K_{\text{tot}}$ such that $k_3 K_{\text{tot}}$ is larger than the maximum value of $v_1([S])$.

Fix $E_{\text{tot}}$ and $K_{\text{tot}}$ satisfying this property. The function $v([S])$ has a last inflection point and decreases towards $L$. Let $-T < 0$ be the slope of the tangent line of $v([S])$ at the inflection point. Since both $v_1$ and $v_2$ depend linearly on $E_{\text{tot}}$, the value of $T$ increases when $E_{\text{tot}}$ and $K_{\text{tot}}$ are increased simultaneously, that is, both are multiplied by the same number larger than 1. Note that the derivative of the first summand of $v([S])$ does not vary.

Therefore, by increasing $E_{\text{tot}}$ and $K_{\text{tot}}$ simultaneously, we can make the slope of the tangent line of $v([S])$ at the inflection point be smaller than $-1$, and the inequalities $k_3 K_{\text{tot}} > L' E_{\text{tot}}$ and $k_3 K_{\text{tot}} > v_1([S])$ still hold. We now proceed to choose $S_{\text{tot}}$ such that the intersection of $v([S])$ with the line $S_{\text{tot}} - [S]$ consists of at least three points, identically to the proof of Theorem 3. This concludes the proof of Theorem 4. □

**Remark 3.** The same observations as in Remark 2 hold for this system as well: $V_{S\to P}$ being of negative type guarantees multistationarity after appropriately choosing $k_3$ such that (2.6) holds.

#### 2.3 Flux through $P$ and $S$

We consider now the system where there is inflow and outflow in the following way

$$0 \underset{k_{\text{outS}}}{\overset{k_{\text{inS}}}{\rightleftharpoons}} S, \qquad P \underset{k_{\text{inP}}}{\overset{k_{\text{outP}}}{\rightleftharpoons}} 0.$$

We allow some of $k_{\text{inS}}$, $k_{\text{outS}}$ and $k_{\text{inP}}$ to be zero, but $k_{\text{outP}}$ must be different from zero. Whether or not the reverse catalysis from $P \xrightarrow{k_h} S$ from Section 2.1 is considered does not change the analysis below. If the reaction is not present, then we simply set $k_h = 0$.

In this case, we have the following result.

**Theorem 5.** *Assume that at least one of $k_{\text{inS}}$ and $k_{\text{inP}}$ is different from zero and that (1.7) holds*

$$\sum_{|I|=n-1} \frac{\sum_{i\in I} c_{I\setminus\{i\},I}}{\sum_{|J|=n-1} \frac{K_J}{K_I}} > \sum_{i\in[n]} c_{[n]\setminus\{i\},[n]}.$$

*Then,*

- *If $k_{\text{outS}}$ is zero, then regardless of the value of $k_{\text{outP}}$, $k_{\text{inS}}$ and $k_{\text{inP}}$, provided one of the last two is non-zero, one can choose $E_{\text{tot}}$ such that the system admits at least two positive steady states.*
- *If $k_{\text{outS}}$ is different from zero, then regardless of the value of $k_{\text{outP}}$ and $k_{\text{outS}}$ one can choose $E_{\text{tot}}$, $k_{\text{inS}}$ and/or $k_{\text{inP}}$ (whichever is non-zero), such that the system has at least three positive steady states.*

*Proof.* By Theorem 2, $V_{S\to P}$ is of negative type. Since the substrate is no longer conserved now, we set the differential equations for $[S]$ and $[P]$ to zero. The sum of the rates of $[S], [P]$ and the concentrations of all intermediate complexes lead to the following relation at steady state:

$$k_{\text{inS}} - k_{\text{outS}}[S] - k_{\text{outP}}[P] + k_{\text{inP}} = 0 \quad \Rightarrow \quad [P] = \tfrac{k_{\text{inS}} + k_{\text{inP}} - k_{\text{outS}}[S]}{k_{\text{outP}}}. \tag{2.10}$$

We set now the differential equation for $[P]$ to zero (as in (2.3) without flux), which gives

$$\begin{aligned} 0 &= \sum_{I\subseteq[n], I\neq\emptyset} \sum_{i\in I} c_{I\setminus\{i\},I}[ES_I] - (k_h + k_{\text{outP}})[P] + k_{\text{inP}} \\ &= \sum_{I\subseteq[n], I\neq\emptyset} \sum_{i\in I} c_{I\setminus\{i\},I}[ES_I] - (k_h + k_{\text{outP}})\tfrac{k_{\text{inS}} + k_{\text{inP}} - k_{\text{outS}}[S]}{k_{\text{outP}}} + k_{\text{inP}}, \end{aligned}$$

which is equivalent to

$$\sum_{I\subseteq[n],I\neq\emptyset}\sum_{i\in I} c_{I\setminus\{i\},I}[ES_I] = \tfrac{(k_h+k_{\text{outP}})(k_{\text{inS}}+k_{\text{inP}})}{k_{\text{outP}}} + k_{\text{inP}} - \tfrac{(k_h+k_{\text{outP}})k_{\text{outS}}[S]}{k_{\text{outP}}}. \tag{2.11}$$

The left-hand side of (2.11) is $v([S]) = V_{S\to P}([S])$ and hence of negative type. We wish now to show that choices can be made such that the line on the right side of (2.11) intersects $v([S])$ in the number of points stated in Theorem 5.

First, assume $k_{\text{outS}}$ is zero, that is, there is no outflow from $S$. Then, the right side of (2.11) is a horizontal line constant at $\frac{(k_h+k_{\text{outP}})(k_{\text{inS}}+k_{\text{inP}})}{k_{\text{outP}}} + k_{\text{inP}}$, which is non-zero provided at least one of $k_{\text{inS}}$ and $k_{\text{inP}}$ is non-zero. Now, we can choose $E_{\text{tot}}$ appropriately such that

$$\max v([S]) > \tfrac{(k_h+k_{\text{outP}})(k_{\text{inS}}+k_{\text{inP}})}{k_{\text{outP}}} + k_{\text{inP}} > E_{\text{tot}}\Big(\sum_{i\in[n]} c_{[n]\setminus\{i\},[n]}\Big).$$

In other words, the horizontal line is above the asymptotic value $L$ in (1.6) of $v([S])$, and below the maximum value that $v([S])$ attains. As a consequence, the horizontal line intersects $v([S])$ in at least two positive points. Hence, since equation (2.11) determines the positive steady states of the network, we conclude that the network has at least 2 positive steady states. This shows the first part of the theorem.

We turn now to the case where $k_{\text{outS}}$ is different from zero and assume (1.7) holds. All we need is to show that values of $E_{\text{tot}}$, $k_{\text{inS}}$ and $k_{\text{inP}}$ can be appropriately chosen such that the line

$$\tfrac{(k_h+k_{\text{outP}})(k_{\text{inS}}+k_{\text{inP}})}{k_{\text{outP}}} + k_{\text{inP}} - \tfrac{(k_h+k_{\text{outP}})k_{\text{outS}}[S]}{k_{\text{outP}}},$$

goes through the last inflection point of $v([S])$ with slope larger than that of the tangent line of $v([S])$ at the inflection point. The slope of this line is $-\frac{(k_h+k_{\text{outP}})k_{\text{outS}}}{k_{\text{outP}}}$. Let $-T < 0$ be the slope of the tangent line of $v([S])$ at the last inflection point. By increasing $E_{\text{tot}}$ if necessary, $T$ can be made larger than $\frac{(k_h+k_{\text{outP}})k_{\text{outS}}}{k_{\text{outP}}}$ (whose value is fixed). Now, the term $\frac{(k_h+k_{\text{outP}})(k_{\text{inS}}+k_{\text{inP}})}{k_{\text{outP}}} + k_{\text{inP}}$ can attain any positive value while varying $k_{\text{inS}}$ and $k_{\text{inP}}$ even if one of the two is imposed to be zero. Changing this value moves the intersection point of the line with the vertical axis. Hence, by choosing $k_{\text{inS}}$ and $k_{\text{inP}}$ appropriately, even if one is imposed to be zero, the line can be made to go through the inflection point.

Repeating the argument in the proof of Theorem 3 in Section 2.1, since the slope of the line is smaller than the slope of the tangent line of $v$ at the inflection point, the two functions must intersect in at least three positive values of $[S]$. This guarantees the existence of at least three positive steady states. Note that using this construction, $[P]$ is positive, as $k_{\text{inS}} + k_{\text{inP}} - k_{\text{outS}}[S] > 0$ (c.f. (2.10)) holds at the intersection points. □

##### 2.4 Positive type curves and multistationarity

In all scenarios covered up to here, we have shown that if $V_{S\to P}$ is of negative type, then multistationarity occurs after appropriately choosing the remaining parameters. The conditions given in Theorem 3, Theorem 4, and Theorem 5 also imply that $V_{S\to P}$ is of negative type.

However, multistationarity is not restricted to $V_{S\to P}$ being of negative type, and multistationarity might occur even if $V_{S\to P}$ is of positive type but has a peak, that is, it is not a monotonically increasing function.

Furthermore, the first part of Theorem 5 guarantees the existence of two positive steady states when $k_{\text{outS}} = 0$, but three steady states might also arise if $V_{S\to P}$ is of positive type with a peak. In this case, the horizontal line in the proof of Theorem 5 can be chosen to intersect $V_{S\to P}$ at three points.

To exemplify this, we consider $n = 2$. With the notation of reaction rate constants given in the main text (Methods section), we choose

$$K_1 = 50,\quad K_2 = 1,\quad k_1 = 1,\quad k_2 = 1,\quad k_3 = 2,\quad k_4 = 1,$$
$$k_5 = 1,\quad k_6 = 1,\quad k_8 = 1/11,\quad k_{10} = 40/256,\quad k_{12} = 1,\quad k_{13} = 1.1,$$

where $K_1 = \frac{k_7+k_8}{k_6}$ and $K_2 = \frac{k_{11}+k_{13}}{k_{10}}$, which define $k_7, k_{11}$. Then $V_{S\to P}$ is of positive type with one peak: it starts increasing, it then decreases below its asymptotic value, and then converges to it from below.

The two panels of Figure S2 show in red the curve $V_{S\to P}$ displaying this behavior, and in blue $V_{P\to S}$. The left panel corresponds to the hydrolytic conversion of $P$ to $S$ studied in Section 2.1, and it shows that after choosing $k_9 = k_h = 0.00021$, $E_{\text{tot}} = 1$ and $S_{\text{tot}} = 5500$, the two curves intersect in three positive points. The right panel corresponds to the inclusion of the fluxes $0 \underset{k_{\text{outS}}}{\overset{k_{\text{inS}}}{\rightleftharpoons}} S$ with reaction rate constants $k_{\text{inS}} = 1, k_{\text{outS}} = 0.143$ and $k_9 = k_h = 0.02$. The two curves intersect in three positive points, and this scenario falls into the first part of Theorem 5 in the sense that $k_{\text{outP}} = 0$.


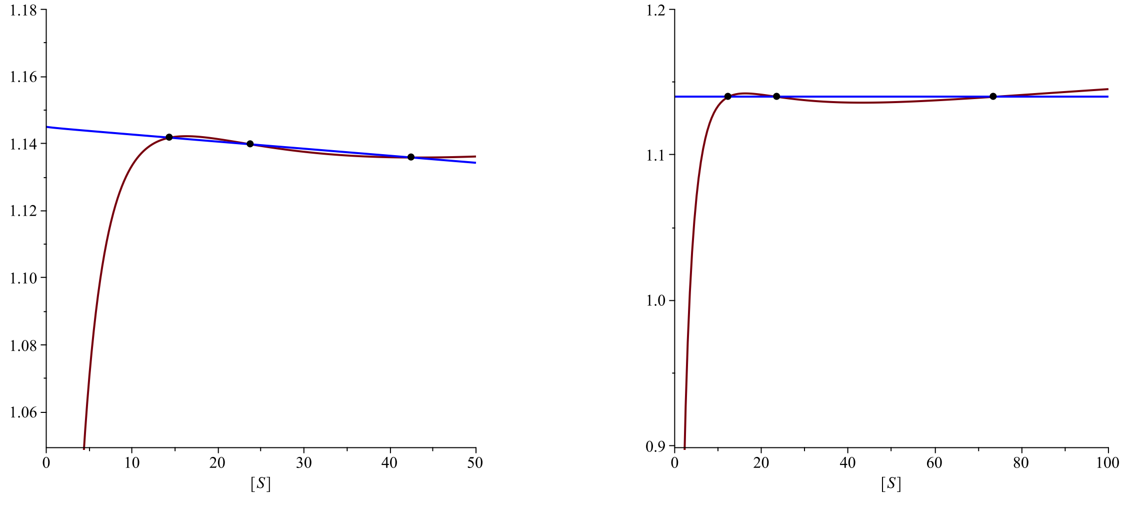


Figure S2: See text.
